# Mid-lateral Cerebellar Purkinje Cells Provide a Cognitive Error Signal When Monkeys Learn a New Visuomotor Association

**DOI:** 10.1101/600221

**Authors:** Naveen Sendhilnathan, Anna E. Ipata, Michael E. Goldberg

**Author notes:** Correspondence to Naveen Sendhilnathan. These authors jointly supervised this work (Anna E. Ipata, Michael E. Goldberg).

## Abstract

How do we learn to establish associations between arbitrary visual cues (like a red light) and movements (like braking the car)? We investigated the neural correlates of visuomotor association learning in the monkey mid-lateral cerebellum. Here we show that, during learning but not when the associations were overlearned, individual Purkinje cells reported the outcome of the monkey’s most recent decision, an error signal, which was independent of changes in hand movement or reaction time. At the population level, Purkinje cells collectively maintained a memory of the most recent decision throughout the entire trial period, updating it after every decision. This error signal decreased as the performance improved. Our results suggest a role of mid-lateral cerebellum in visuomotor associative learning and provide evidence that cerebellum could be a generalized learning system, essential in non-motor learning as well as motor learning.

---

The cerebellum is a learning machine: it uses errors in prior performance to shape behavior for future performance (*1-4*). For over 150 years, the cerebellum has been studied mostly for its role in the regulation of movement (*2, 5, 6*) and motor learning (*7*). Much evidence shows that the cerebellum facilitates motor learning using prior motor errors to adjust the parameters of upcoming movements(*1, 2, 8-13*). However, several recent clinical(*14-18*), imaging(*19*) and anatomical(*20, 21*) studies have suggested that cerebellar processing goes beyond the realm of motor control. Recently, a consensus has been established that the cerebellum, basal ganglia and the prefrontal cortex are reciprocally connected and are a part of a distributed cortical network(*22, 23*) suggesting that the cerebellum and the prefrontal cortex may work in tandem in generating complex behavior beyond that of fine-tuning movement. This anatomical connection thus raises an important functional question: does the cerebellum contribute to non-motor learning(*24-26*) ?

Here we asked if mid-lateral cerebellum provided a cognitive error signal that could facilitate the learning of an arbitrary visuomotor association. During visuomotor association learning, animals use information available in the outcome of the recent decision to learn that a given visual stimulus is associated with a given movement and report their choice through well-learned movement that does not necessarily change through the paradigm. Hence, this paradigm provides a convenient way to study the effects of higher order processing, decoupled from motor learning, in a timescale that is experimentally tractable. Learning a novel visuomotor association involves understanding that the presence of an arbitrary symbol instructs the subject to make a particular movement even though there is nothing about that symbol that describes the movement it instructs, as a red traffic light instructs a driver to press the brake. In this study, we show that the Purkinje cells (P-cells) in the midlateral cerebellum track the learning of a new visuomotor association by reporting the outcome of the monkey’s most recent decision, a error signal that occurred when the monkey was learning new visuomotor associations but not when the monkey had already learned the task. The error signal is cognitive in nature in that it does not describe a mistake in the parameters of the movement, but rather it describes the consequence of failing to make the association.

## Cerebellum and visuomotor associative learning

We trained two monkeys to perform a two-alternative forced-choice discrimination task, where the monkeys associated one of two visual symbols with a left-hand movement and the other symbol with a right-hand movement. The monkeys began each trial by placing their hands on each of two bars, after which one of the two symbols appeared on the screen and the monkeys lifted the hand (arbitrarily) associated with that symbol to earn a liquid reward (**Fig 1A**). The liquid reward and a beep paired with opening of the solenoid were delivered immediately (with a delay of 1 ms) after the initiation of the correct hand movement. We began each recording session began with an overtrained association, using the same familiar symbols and visuomotor associations. We then changed the symbols to a pair of novel fractal stimuli that the monkeys had never seen before and that were different each session. The monkeys had to learn the arbitrarily assigned correct symbol-hand association through trial and error, which they did usually in ∼50-70 trials on an average through an adaptive learning mechanism (**Fig 1B**). The monkeys typically used a win-stay-lose-switch strategy in the learning phase (**Fig 1B** inset).

**Figure 1:**
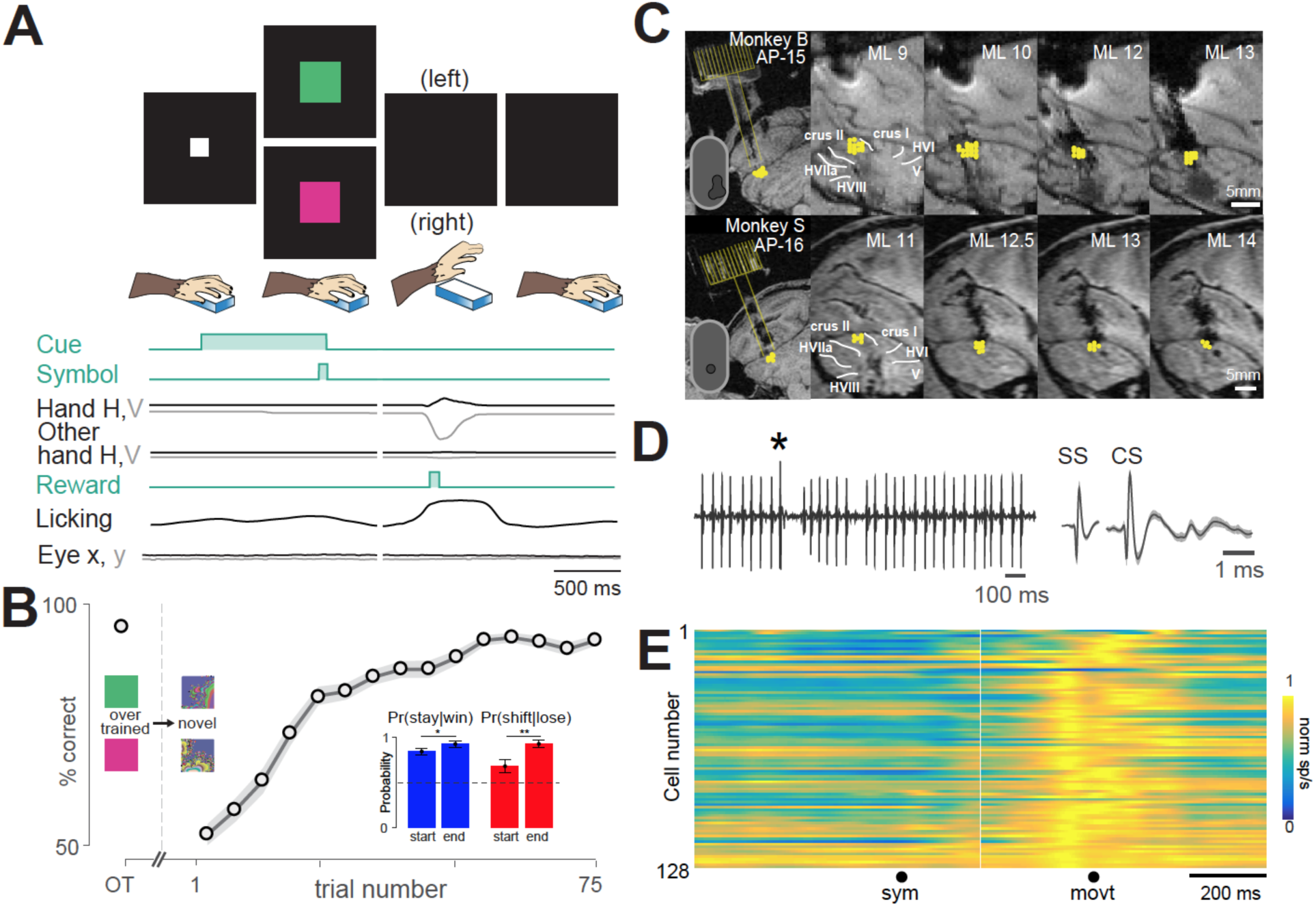
Mid-lateral cerebellum and visuomotor association. **A**. Two-alternative forced-choice discrimination task (top) and parameters (bottom): The monkeys pressed each bar with a hand, and a white square appeared that served as the cue for the start of the trial. Then one of the two visual symbols appeared briefly. The monkey lifted the hand associated with the presented symbol to earn a reward immediate. The correct symbol-choice association is shown for the overtrained task (Green symbol – left hand; Pink symbol – right hand). **B**. Mean learning curve of both monkeys while they learned a new visuomotor association from an overtrained (OT) one. Inset shows the strategy of learning at the start and the end of learning. The horizontal broken line is the chance level. **C**.T1 MRI of the cerebellum recording locations (yellow markers) in both monkeys. The first panel is the coronal slice showing the chamber location and the burr-hole tract (yellow). Inset shows the chamber and the burr-hole location from bird’s eye view. The next four panels are sagittal reconstructions showing recording locations (yellow markers). **D**. Representative recording from a mid-lateral Purkinje neuron showing simple spikes (SS) and complex spike (CS; marked by *). Each row shows the trial-averaged spike response of a single P-cell aligned on the symbol onset (sym) and bar-release hand movement onset (movt).

We recorded 128 P-cells near crus I and II of the mid-lateral cerebellum of two monkeys (**Fig 1C-D**; see methods; 96 from monkey B; and 32 from monkey S). Similar to previous observations in the mid-lateral cerebellum(*27*), during the overtrained condition, most P-cells (106/128) significantly increased their firing rate during the bar-release hand movement (**Fig 1E**). The movement related-cells responded similarly during right and left hand movements, and responded similarly to both symbols (**Fig S1)**. The remaining 22 neurons did not show any significant movement-related activity.

Given the prominent role of the cerebellum in motor control(*2, 5*), if these P-cells were purely motor neurons, we would expect the neural activity to not change when the monkeys had to learn a new visuomotor association, as long as the hand movement made by the monkeys remained the same. However, contrary to this, we found that when the task changed from the overtrained to the novel condition (**Fig 2A**), the activity of most P-cells (both selective and non-selective for bar-release hand movement, see **Supplementary Table 1**) changed dramatically (N = 105, *P* = 10^−21^; KS test) in several intervals of the trial (for single cell example, see **Fig 2B**). Across the population, the change in activity occurred at different times for different cells.

**Figure 2:**
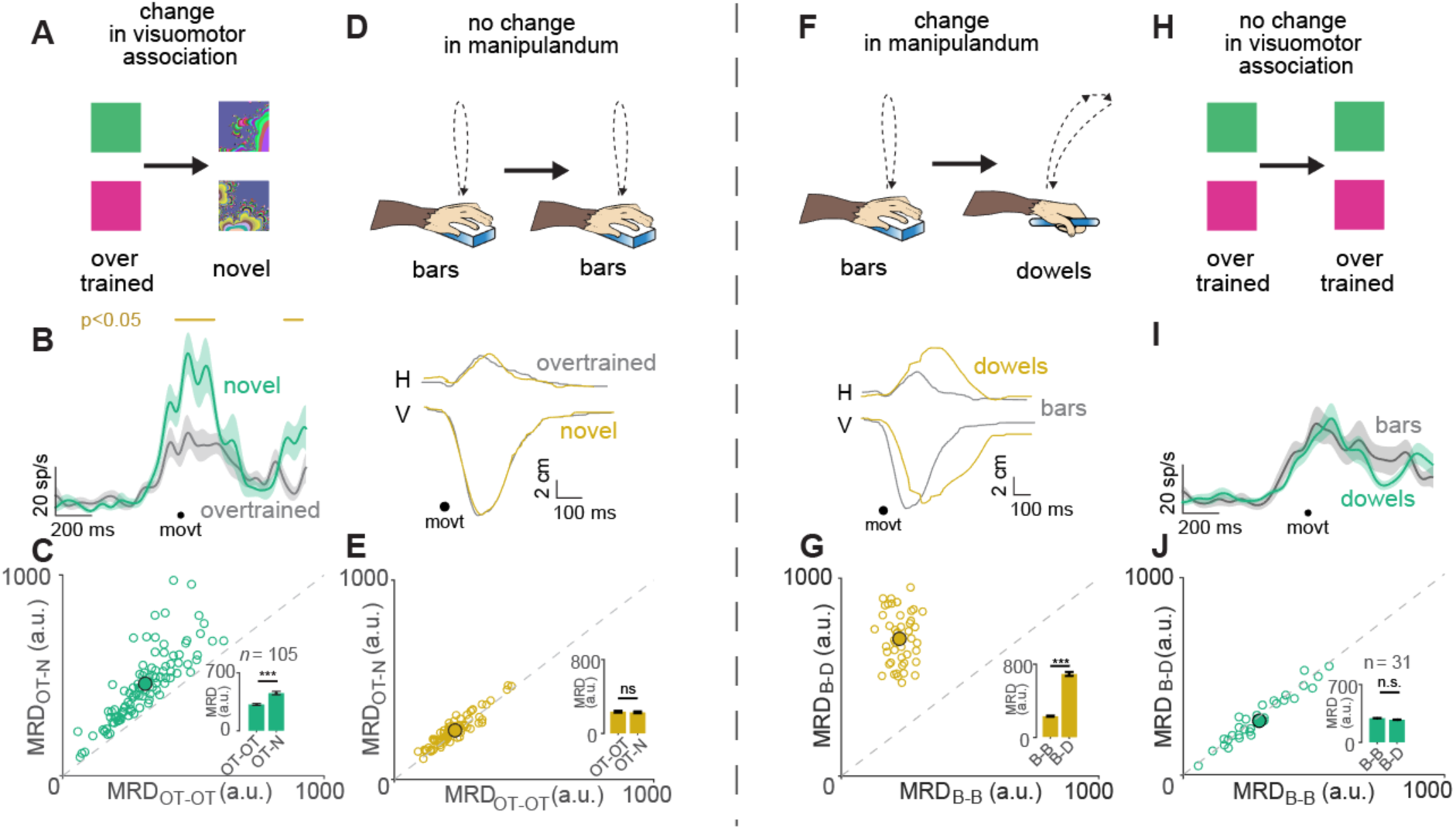
P-cells responded to change in visuomotor association learning and did not encode specific motor kinematics. **A**. Task in which there was a change in cognitive load: visuomotor association changed from overtrained to novel association. **B**. Spike density plot of a representative P-cell aligned to bar-release hand movement onset (movt) that responded to this change in cognitive load by firing differently between the overtrained (gray) and novel (green) conditions. The epochs where this difference was significant (p<0.05, t-test) is shown by a gold line on the top. **C** Scatter plot of Mean RMS Distance (MRD) for within overtrained (OT) condition (OT-OT) vs MRD for across overtrained and novel (N) condition (OT-N), obtained from method described in **Fig S2**. Each open circle is a P-cell and the mean value is shown as a filled circle. The inset shows the mean MRD for OT-N and OT-OT conditions. *** means *P* = 10^−8^ Mann-Whitney U-test. **D**. A cartoon bar-release hand movement trajectories (top) and the actual movement trajectories decomposed into horizontal (H) and vertical (V) components traces (bottom) for overtrained (grey) and novel (yellow) conditions. **E**. Same as 1c but for bar-release hand movement trajectories. n.s means *P* = 0.3822, paired t-test **F**. Top: A cartoon showing different hand movement trajectories with the change in manipulanda from bars (B) to dowels (D). Bottom: Actual movement trajectories decomposed into horizontal (H) and vertical (V) components traces for bars (grey) and dowels (yellow) conditions. **G**. Same as 2e but for hand movement trajectories in bars-dowels condition. *** means *P* = 10^−32^, paired t-test **H**. Task in which the cognitive load didn’t change: visuomotor association remained in the overtrained condition. **I**. Same representative neuron from 1b when the movement changed but the association did not. **J**. Same as 1c but for neural activity in bars-dowels condition. n.s means *P* = 0.2330, paired t-test

To show that the activity profile between the two conditions was significantly different despite this heterogeneity, in a way that is neither epoch nor neuron (statistical distribution) dependent, we computed the root mean squared (rms) distance between the spike density functions across the overtrained and novel conditions with repeated random sampling and compared the resulting distribution’s mean (MRD_across_) with the mean (MRD_within_) from a distribution of two random samples repeatedly drawn without replacement, from within the overtrained condition, compensating for differences due to reaction times (**Fig S2;** see methods). The within-condition MRD values, for the population, were significantly lower than the across-condition MRD values indicating that the change in neural activity in the novel condition was significantly different from the activity in the overtrained condition (*P* = 10^−8^ Mann-Whitney U-test; **Fig 2C**).

Although the neural activity changed dramatically at the symbol switch, by monitoring the gross hand movement of the monkeys (see methods), we verified that the monkeys showed no significant difference in motor kinematics (**Fig. 2D-E,** *P* = 0.3822, paired t-test. Also see: **Fig S3 A-D**). Finally, a change in reaction time at the symbol switch did not contribute to the change in neural activity either: the reaction time did not change at the symbol switch on 24/105 (23%) of sessions even though the monkeys’ performance decreased significantly but nevertheless, the neural activity changed significantly in these sessions (**Fig S4**). This also indicates that the change in neural activity was independent of motor learning.

If the neurons were truly hand-movement invariant and only changed their activity when the monkeys had to learn a new visuomotor association, then we should expect the neural activity to remain unchanged in a condition where the monkeys reported their choices of the same visuomotor associations with different hand movements. To test this possibility, we changed the manipulanda, which forced a change in the movements associated with manipulandum release (**Fig 2F**) while keeping the visuomotor association the same (**Fig 2H**). We started certain sessions with the bars, but on a randomly chosen trial, we switched the manipulanda to a pair of dowels (cylindrical rods), on which the monkeys were seldom trained for a long time. Since there was no new visuomotor associative learning, the monkeys performed close to 100% in both conditions (P = 0.7895; t-test) and the reaction times were comparable (P = 0.7696; t-test). Although the kinematics of the movement changed markedly (**Fig. 2G,** *P* = 10^−32^, paired t-test Also see: **Fig S3 E-H**), the neural activity did not change significantly (**Fig 2 I-J;** *P* = 0.2330, paired t-test). These results taken together, suggest that P-cells encoded changes in association learning but not changes in motor kinematics which is a major departure from our current understanding of the cerebellum’s role in motor control.

## Learning Contingent Error Signal for Performance Monitoring

In the overtrained task, the monkeys almost always performed the task correctly (**Fig 1B**). However, after the symbol switch, during early learning, the monkeys had to use their success or failure on every trial to learn the correct association. Based on the win-stay-lose-switch strategy used by the monkeys during learning (**Fig 1B inset**), we hypothesized that the P-cells must have information about the one-back decision during learning. Therefore, we compared the trial-averaged spike density functions for correct and wrong outcomes, from soon after the reward delivery (or its absence) on one trial through the reward delivery (or its absence) in the next trial to see if there are any differences. We did this by calculating the root mean squared distance between the trials with real vs shuffled outcome labels by iterative random sampling. We found that the same 105 cells that showed a significant modulation in activity at the symbol switch also showed a significant difference in activity between correct and wrong outcomes during learning (P = 2.359e-4, ranksum test). By repeating this analysis on N-back (up to 3) and N-forward (up to 2), we found that the P-cells only reported one-back outcome, which is consistent with the behavior (**Fig 3A-B**; P values: 3-N = 0.6411; 2-N = 0.8025; 1-N = 2.359e-4; 1+N = 0.2680; 2+N = 0.9574, ranksum test; this is discussed further in the next section). Additionally, the cells that did not have any information about the trial outcome in 1-N condition did not have any information about the trial outcome in N-3, N-2, 1+N or 2+N trials as well (**Fig 3A-B**). We restrict our further analyses to only those cells that showed a statistically significant change between the correct and wrong outcomes in the 1-N case, during learning.

**Figure 3:**
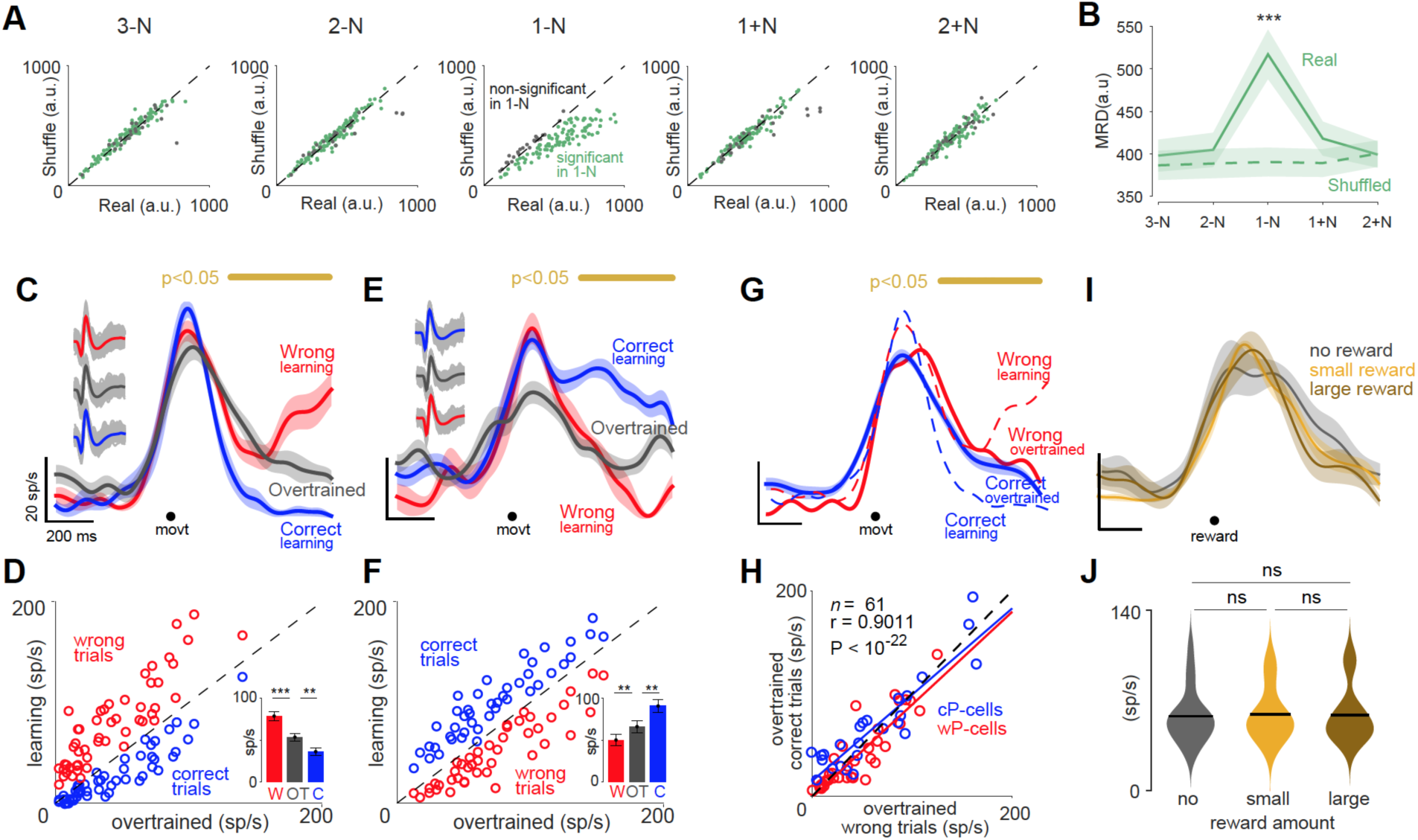
P-cells reported one-back decision’s outcome during learning but not when overlearned. A. RMS distance between spike density functions of real vs shuffled correct and wrong trials for all the recorded neurons. Green: P-cells that showed significant difference (P>0.05; ranksum test) in the 1-N condition; Black: P-cells that did not show a significant difference (P<0.05; ranksum test) in the 1-N condition. B. MRD (mean rms distance) from all the cells that showed a significant difference in a, in the real (solid line) and shuffled (broken line) case for all the five conditions; Only the 1-N condition was significant (P values: 3-N = 0.6411; 2-N = 0.8025; 1-N = 2.359e-4; 1+N = 0.2680; 2+N = 0.9574; ranksum test). C. Spike density function of a representative wP-cell for wrong trials during learning (red), correct trials during learning (blue) and all trials in the overtrained condition (grey) aligned to movement onset. The gold bar on the top indicates the continuous time-period when the activity in the wrong trials and the correct trials were significantly different from each other (P<0.05; t-test. The inset on the left shows the spike waveforms for all the simple spikes isolated for three conditions. D. Mean neural activity in the delta epoch of all trials in the overtrained condition (before; abscissa) plotted against the mean neural activity in the delta epoch of correct trials during learning (blue) and wrong trials during learning (red) for wP-cells. Broken diagonal line is the line of unity. (wP-cells: W-OT: P = 10^−6^, ranksum test; OT-C: P = 0.0025, ranksum test) E. Same as a, but for a representative cP-cell. F. Same as b, but for all cP-cells (W-OT: P = 0.0034, ranksum test; OT-C: P = 0.0021, ranksum test). G. Spike density functions of the same wP-cell from 3a; but for wrong trials during learning (red broken), correct trials during learning (blue broken), wrong trials in the overtrained condition (red solid) and correct trials in the overtrained condition (blue solid) aligned to movement onset. H. Activity in the wrong trials of the overtrained condition plotted against the activity in the correct trials of the overtrained condition in the delta epoch, for wP-cells (red) and cP-cells (blue). I. Spike density functions for no-reward (grey), small reward (gold) and large reward (brown) trials in the overtrained condition, aligned to movement onset. J. A violin plot showing the activity across the population of n = 25 neurons in the three trial types for the overtrained condition. Horizontal black line indicates the mean for each group. ns means not significant (no-small: P = 0.9125; small-large: P = 0.8605; no-large: P= 0.9898 Mann-Whitney U-test).

Within this group, we found two populations of P-cells that reported the outcome of the most recent decision during learning (see methods). After a wrong outcome, wrong-reporting Purkinje cells (wP-cells; *N*=54; **Fig 3C**) increased their firing rate significantly more relative to their activity in the overtrained condition (P = 10^−5^ t-test; **Fig 3D**). After a correct outcome, their firing rate was significantly lower relative to their activity in the overtrained condition (P = 0.0021 t-test; **Fig 3D**). In contrast, the firing rate of correct-reporting Purkinje cells (cP-cells *N*=51; **Fig 3E**) was significantly greater or lesser than their firing rate respectively, after a recent correct (P = 0.0442 t-test) or wrong outcome (P = 10^−4^ t-test), relative to the activity in the overtrained condition (**Fig 3F**). Although the P-cells changed their activity profile from the OT to the N condition in several intervals of the trial (**Fig 2**) during learning, this difference in activity after correct and wrong outcomes did not occur throughout the trial but occurred only in brief epochs (see next section). We performed a t-test between the neural activities following correct and wrong outcomes, during learning, for each neuron separately and identified the continuous time period when this activity difference was significant (P<0.05) and referred to it as the delta epoch (see methods). For the population of P-cells, the delta epoch lasted for 256 ± 25 ms and had information about the outcome of the monkey’s most recent decision. We hypothesized that this activity in the delta epoch provided a learning contingent error signal that monitored the monkeys’ performance as they learned a new visuomotor association.

The P-cells reported the outcome of the decision only during learning and not during the overtrained task: both the correct and the occasional wrong trials in the overtrained condition had similar neural activities (**Fig 3G-H**). This is despite the fact that the granule cells one synapse upstream to the P-cells have information about the delivery or omission of reward in the absence of learning(*28*). If the P-cells did not use the reward information during the overtrained condition, then we should expect the P-cell activity to be similar in rewarded or reward omitted correct trials in the overtrained condition. To test this, we randomly chose a reward probability for a given trial as no, small or large reward. The neurons did not show any significant changes in activity among these conditions, indicating that the P-cells were not sensitive to the amount of reward or its absence in the overtrained condition (**Fig 3I-J**). Therefore, P-cells reported the trial outcome only when relevant to learning. This further indicates that the cerebellar cortex is essential only for learning while the consolidation of the learned associations maybe occur elsewhere in the brain(*29*).

Furthermore, although granule cells (*28*) and climbing fiber driven complex spikes (*30*) have reward expectation signals, the simple spikes of mid-lateral P-cells did not. To demonstrate this, we trained the monkeys on a stimulus-reward association task (see methods and **Fig S5A**) where the monkeys were rewarded 800 ms after the presentation of a small red square, a task that required neither learning nor even a response on the monkey’s part. The monkeys did not make any hand movement during this task. Although the monkeys anticipated the reward by licking the juice spout with an increased rate during the trial, in a manner not different from anticipatory licking in the overtrained visuomotor association task, the P-cells did not respond in this simple stimulus-reward association task (**S5B-C;** P = 10^−7^, Mann-Whitney U-test). Taken together, these results suggest that the neural activity that started to increase before the reward period in the overtrained condition was task dependent (hand-movement related) and represented neither reward anticipation nor licking. Finally, the dissociation between reward and neural activity during the overtrained and reward expectation tasks suggests that the activity during learning did not result from sensory and motor events associated with the reward, such as the click of the solenoid or the monkey’s licking or swallowing of the liquid reward.

## P-cells encoded one-back memory in bursts

Although the population of P-cells fired precisely at the same time in response to the task-related bar-release hand movement (**Fig 1E**), different P-cells had delta epochs at different times, sprinkled throughout the trial period. The earliest time that any P-cell reported the outcome of the immediately preceding decision was 310 ms for wP-cells, and 330 ms for cP-cells after the reward onset. This provided an approximate estimate of the reward information latency (RIL; the time from reward delivery to neural response) for mid-lateral P-cells. From the RIL of a given trial (n) through the RIL of the next trial (n+1), the P-cells only reported the most recent, that is the one-back decision’s outcome (1-N outcome; see **Fig 4A**; also see **Fig 3A**), however with different latencies. That is, some neurons reported the outcome immediately, with short latency and hence had delta epochs during the post-reward epoch (in the inter trial interval, after the RIL; **Fig 4B-C leftmost**), but others reported the same outcome with longer latencies so that their delta epochs carried over to the next trial (**Fig 4B-C left-center to rightmost**), but all of them still reported the one-back decision (1-N decision). Therefore, although the delta epochs were themselves short (∼250 ms), the delta epochs of all the P-cells combined, collectively tiled the entire trial interval (**Fig 4E**).

**Figure 4:**
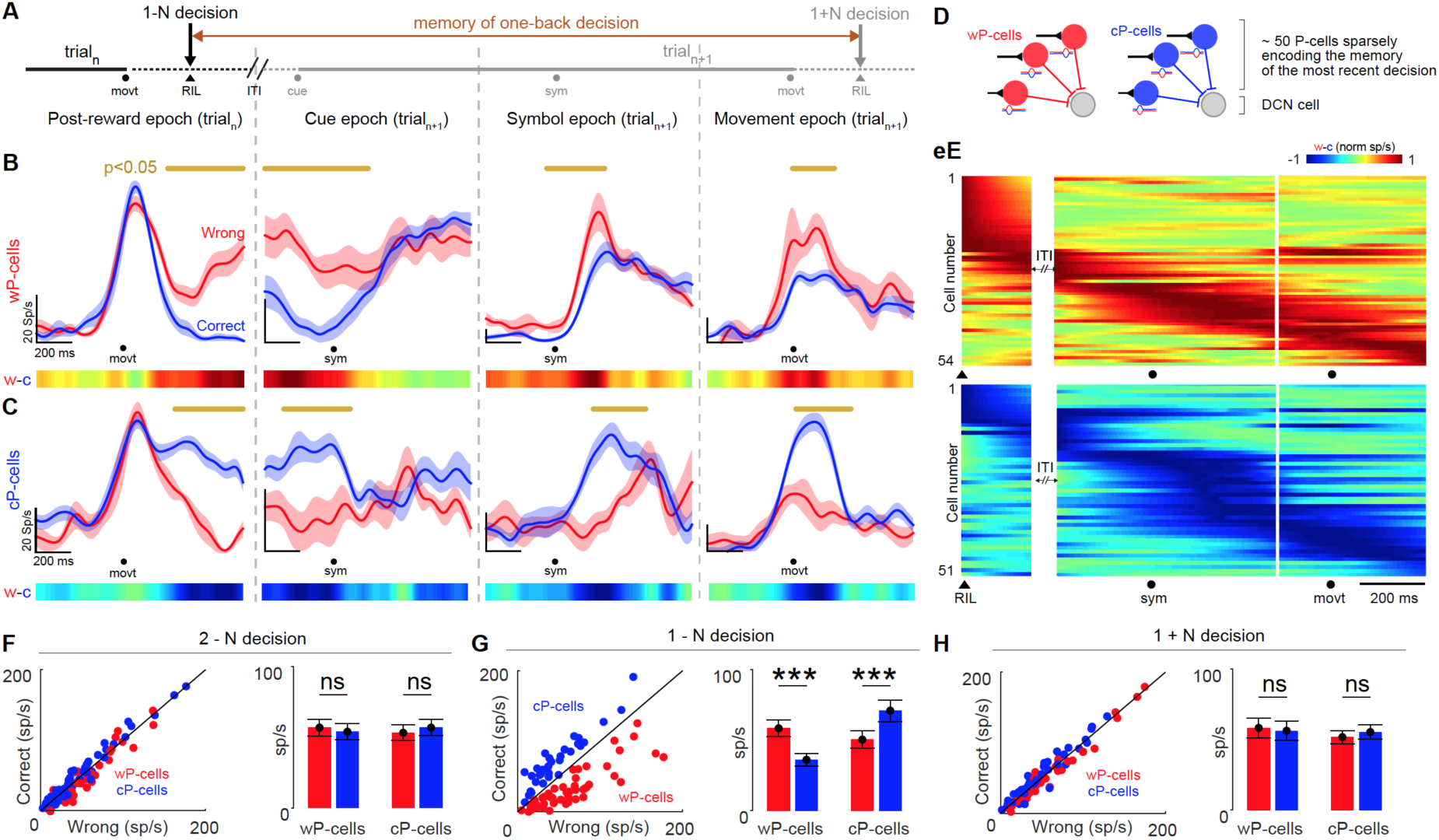
P-cells reported the outcome of the one-back decision in continuous small epochs that tiled the entire trial period. A. A schematic illustration of trial structure with two consecutive trials (solid lines) separated by ITI (Inter-trial interval; broken line) highlighting various epochs. RIL: reward information latency, the time taken for the reward information to reach the P-cell after reward delivery. From one RIL to the next RIL, the P-cells maintain the memory of the most recent, one-back, decision as explained below. B. Representative wP-cells whose activities were higher after a wrong trial (red) relative to a correct trial (blue). The top gold line indicates the time when the difference in activity was continuously significant (P<0.05 t-test). The heat map in the bottom indicates the difference between wrong and correct traces. Leftmost neuron is aligned to movement onset with the delta epoch after RIL. The left-center neuron is aligned to symbol onset of next trial with the delta epoch in the cue epoch. The right-center neuron is aligned to symbol onset of next trial with the delta epoch in the symbol epoch. The rightmost neuron is aligned to movement onset of the next trial with the delta epoch in the movement epoch. Note that the reward was delivered 1 ms after the correct movement onset. C. Representative cP-cells whose activities were higher after a correct trial (blue) relative to a wrong trial (red). Same convention as b. D. Hypothesized anatomical connection between P-cells and downstream deep cerebellar nuclear cells (DCN). E. Each row shows the difference in neural activities between trials following wrong and correct trials of a single P-cell aligned on the symbol onset (sym) and bar-release hand movement onset (movt) arranged in the increasing order of the peak differences for wP-cells (top) and cP-cells (bottom). RIL: Reward Information Latency; ITI: Inter-trial interval. F. Memory of two back decision (2-N): Left: Trial averaged neural activity after 2-N wrong trials (abscissa) versus 2-N correct trials (ordinate) in the delta epoch for wP-cells (red markers) and cP-cells (blue markers). Diagonal line is the line of unity. Right: The mean activities of the population of wP-cells for 2-N wrong (red) and 2-N correct (blue) trials were not significantly different (P = 0.1068, ranksum test) and nor were the mean activity of the population of cP-cells for 2-N wrong (red) and 2-N correct (blue) trials (P = 0.5397, ranksum test). G. Memory of one back decision (1-N): Left: Same as above, but for 1-N wrong (abscissa) and 1-N correct trials (ordinate). Right: The mean activities of the population of wP-cells for 1-N wrong (red) and 1-N correct (blue) trials were significantly different (P = 5.41e-4, ranksum test) and so were the mean activity of the population of cP-cells for 1-N wrong (red) and 1-N correct (blue) trials (P = 1.1862e-5, ranksum test). H. Prediction of impending decision (1+N): Left: Same as above, but for 1+N wrong (abscissa) and 1+N correct trials (ordinate). Right: The mean activities of the population of wP-cells for 1+N wrong (red) and 1+N correct (blue) trials were not significantly different (P = 0.2333, ranksum test) and nor were the mean activity of the population of cP-cells for 1+N wrong (red) and1+N correct (blue) trials (P = 0.0711, ranksum test).

As we showed already (**Fig 3A**), the P-cell activity in the delta epoch did not predict the impending decision’s outcome (1 + N) nor did it have a memory of the history of previous decision (2 - N) before the most recent decision (1 - N) (see methods, **Fig 4F-H, Fig S6-7**). That is, the population of P-cells collectively maintained a memory of only the most recent decision from one RIL to the next, updating it after every RIL during learning (**Fig 4E**).

## Properties of the Error Signal

Two lines of converging evidence suggest that the delta epoch was an intrinsic property of each P-cell and did not arise due to random chance or threshold crossing in a population of neurons with similar properties. First, we shuffled the correct and wrong labels for all the trials during learning, and we found that the delta epoch did not emerge for any P-cell (**Fig S8**). Second, on some sessions (*n* = 24), we reversed the symbol-hand associations once the monkeys had learned the novel associations (see methods for criterion for ‘learned’). For a given neuron, the delta epoch almost always occurred at the same time, reporting the same type of decision, in the novel and the reversal learning conditions (**Fig S9**; least square slope = 0.9831; rho = 0.9709, P< 10^−10^, Pearson correlation). Thus, a given P-cell consistently showed the delta epoch at the same time in every trial during learning.

We have at least five lines of evidence suggesting that the delta epoch did not occur due to sensorimotor parameters. First, the neural activity was not selective for the presented visual symbol (**Fig S10A-B**) or the hand that the monkey used to report the choice (**Fig S10C-D**); but nevertheless, differed between wrong and correct trials. Second, the activity in the delta epoch was independent of manual reaction time: we analyzed the trial-by-trial reaction time and its correlation with neural activity for wrong trials and correct trials separately and we found no difference (**Fig S10E-G**). Third, during overtrained or during learning conditions, neither the hand movement trajectories of the hand used to report a decision (OT: P = 0.2724, ranksum test; learning: P = 0.3474, ranksum test; **Fig S10H**), or the hand movement trajectories of the other hand (OT: P = 0.9732, ranksum test; learning: P = 0.9274, ranksum test; **Fig S10I**) nor the time at which the monkeys returned their lifted hand back to touch the bar (OT: P = 0.3663, ranksum test; learning: P = 0.1452, ranksum test; **Fig S10J**) were significantly different between the correct and wrong trials during learning nor during the overtrained condition and therefore, the differences in the movements of both hands could not have contributed to the differences in firing of the neurons in the delta epoch. Fourth, since the monkeys were free to move their eyes with no constraints whatsoever, the non-task related eye movements made by the monkeys were highly variable across sessions and conditions although they mostly fixated at the cue and symbol. The average eye movements following correct and wrong trials were not different from each other (P = 0.7703, ranksum test) and thus, consistent neural activity despite variable eye movements suggests a dissociation between the two (**Fig S10K-M)**. Finally, the dissociation between the reward and neural activity during the overtrained and reward expectation tasks suggests that the activity during learning was not affected by the sensory and motor events associated with the reward, such as the click of the solenoid or the monkey’s licking or swallowing of the liquid reward. In the overtrained condition, the monkeys licked more for correct trials than for wrong trials after making a decision (with a hand movement) (**Fig S10N**) while the neural activity was not different between the two trial types. In the novel condition, during early learning, they licked more after they made either decision (with a hand movement) anticipating a reward regardless of the task outcome and hence their licking behavior looked similar for correct and wrong trials (**Fig S10N**) although the neural activity differed between two trials types. These results thus strongly suggest that the neural activity in the delta epoch was independent of changes in licking behavior or a sensory memory of the solenoid click. Nevertheless, we cannot exclude the possibility that other body movements unaccounted for by our tracking and analyses, including minor differences in grip, movement of digits, minor postural adjustments etc., could potentially contribute to the observed changes in neural activity.

## The Error Signal Decreased with Learning

Finally, we investigated how this transient change in activity during initial learning changed through the time course of learning. An error signal that that could be used in learning should be maximal when the learning begins, and approach zero as the learning completes. We analyzed changes in neural activity in the delta epoch, through learning until the monkeys learned the associations: as would be expected for a learning contingent error signal, the difference between the activity following a correct trial and wrong trial in the delta epoch monotonically decreased significantly for both types of neurons, as the monkeys learned the association (**Fig 5A**). Importantly, only the activity in the delta epoch, but not any other randomly chosen epoch, provided the error signal that changed systematically with learning (**Fig 5B**).

**Figure 5:**
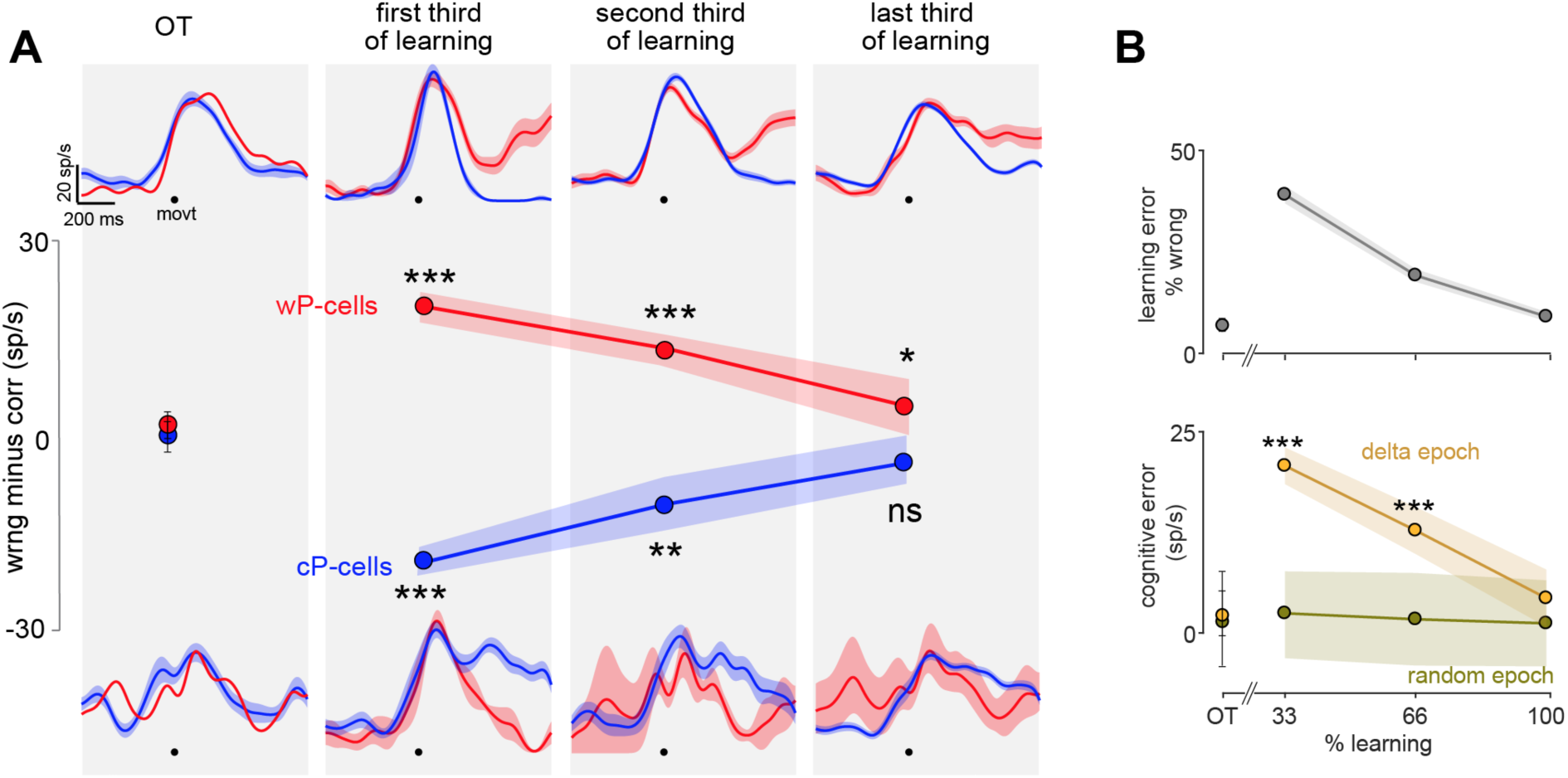
P-cell activity tracked learning. A. Difference between wrong and correct trials before learning (OT), in the first, middle and last 33% of learning for wP-cells (red; first *** means P = 4.3e-5; second *** means 0.0021; * means P = 0.0414) and cP-cells (blue; first *** means P = 0.0091; second *** means 0.0111; n.s means P = 0.0633). B. Top: average learning error from all sessions (grey). Bottom: Average magnitude of the error (| wrong – correct |) calculated in the delta epoch (yellow) and from a random sample of 200 epochs per neuron (green) for all neurons as a function of learning. The shaded region in the delta epoch case is s.e.m. But the shaded region in the random epoch condition is 2 x stdev.

A reinforcement learning framework combined with a drift diffusion model sufficiently explains our data. Here, we modelled the action selection, in response to a presented symbol, through a race between two action choice accumulators (left and right hand release) that drifted until one of them reached a threshold. We then hypothesized the trial-to-trial changes in the rate of the accumulators to be given by the P-cell activity in the delta epoch that changed with learning (**Fig S11**).

## Discussion

Given there is an anatomical substrate for cerebellar participation in non-motor processes, we show for the first time that the cerebellum actually provides a neural correlate of visuomotor association learning. When monkeys have to learn a new visuomotor association, P-cells significantly changed their activity profile and as monkeys learned by trial and error, P-cells collectively maintained a memory of outcome of the monkey’s most recent decision, a learning contingent cognitive error signal that decreased with improving performance.

Two distinct types of cerebellar error signals have been previously discussed in the literature: first, a simple spike rate signal that encodes the kinematic error of an effector and is used for feedback control of that effector(*31-33*), and second, a complex spike signal that reflects an error that is used to change the synaptic weights of the granule cell input to the P-cells(*7, 34*). The error signal that we report here is similar to the first type, in that the errors are encoded in simple spike output, but with the critical difference that the simple spike output is not linked to the motor plant, but rather signals task-dependent, non-sensorimotor errors that are likely used by higher brain areas for learning. Furthermore, it is unlikely that a trial-by-trial difference in the simple spike rate of more than 30 sp/s in the delta epoch could be caused solely by synaptic depression caused by complex spike input(*7, 34*). Therefore, it might not be the case that the same P-cells we report also use the second type of error signal to shape the first type, over learning. In keeping with this, we did not see a similar tiling effect of the complex spike signals across the population. These observations suggest that the error signals may be inherited from mossy fiber inputs, such as those relaying information from cerebral cortex, which is known to contain higher-level cognitive signals(*22, 28*)

The information about the reward resulting from a particular action is often revealed only after a delay (RIL), while in the meantime the animal may carry out several other actions before collecting the reward that specifically resulted from a previous action. Therefore, making associations between a particular action and its corresponding outcome correctly might pose a problem. This is referred to as the problem of temporal credit assignment (*35*). One solution to this problem is by using short term memory signals, called eligibility traces, related to the selected actions, that can facilitate action-outcome association even when the outcome is delayed. Because ∼50 P-cells synapse onto a single dentate nucleus neuron(*36*), it is possible that the bits of information carried by the P-cells in the delta epoch are integrated by the dentate neurons downstream, which can then provide an eligibility trace signal to the caudate nucleus and prefrontal cortex to enable action selection. This is supported by our model, which is in accordance with a reinforcement learning framework.

The cerebellum can thus function at two different levels: at one level, the cerebellum can send a motor error signal to the motor areas of the brain to regulate the motor kinematics of muscles and at another level, send a cognitive error signal to the cognitive areas of the brain to regulate the cognitive parameters during a learning process(*24*). Although we are currently unaware if these signals are necessary for cognitive learning, in the absence of such cerebellar cognitive error signals, one could hypothesize that the brain would have to perform cognitive learning less rapidly and less optimally.

## Acknowledgments

We thank Glen Duncan for highly creative and wonderful electronic assistance, John Caban for designing and building the manipulandum changer and the video mount and all other creative machining, Matthew Hasday for superb machining, Dr. Girma Asfaw, Dr. Moshe Shalev for animal care, Dr. Mulugeta Semework for assistance with systems programming, MRI and surgeries, Dr. Vincent Prevosto for helping us install the CED1401 and interfacing it with REX, and Lisa Kennelly, Whitney Thomas and Holly Cline for facilitating everything.

## Data and materials availability

All data is available in the main text or the supplementary materials. Raw data and codes are available upon reasonable request.

## Supplementary Materials

### Animal subjects and surgery

We used two male adult rhesus monkeys (*Macaca mulatta*), B and S, weighing 10-11 kg each, for the experiments. All experimental protocols were approved by the Animal Care and Use Committees at Columbia University and the New York State Psychiatric Institute, and complied with the guidelines established by the Public Health Service Guide for the Care and Use of Laboratory Animals. We located the cerebellum in each monkey using Siemens (at the Zuckerman Institute) or GE (at the NYSPI) 3T magnet, with a tungsten recording electrode placed at a position from which we had recorded hand-movement related activity with changes at the symbol switch. Using standard sterile surgical techniques and endotracheal isoflurane general anesthesia, we implanted 10–15 titanium screws in the monkeys’ skull and used them to anchor an acrylic cap in which we placed a head-fixing device and the recording chamber. We left the bone intact and made 3 mm burr holes through which we then could insert the electrodes. We used two recording cylinders, on the left hemisphere of each monkey.

### Task

We used the NIH REX system for behavioral control(*37*). The monkey sat inside a dimly lit recording booth, with its head firmly fixed, in a Crist primate chair 57 mm in front of a back-projection screen upon which visual images were projected by a Hitachi CP-X275 LCD projector controlled by a Dell PC running the NIH VEX graphic system.

#### Two-alternative forced-choice discrimination task

The task began with the monkeys grasping two bars, one with each hand, after which a white 1° x 1° square appeared as a trial cue for 800 ms. Then one of a pair of symbols appeared, briefly for 100 ms in some sessions or until the monkey initiated a hand response in other sessions, at the center of gaze. One symbol signaled the monkey to release the left bar and the other to release the right bar. We rewarded the monkeys with a drop of juice for releasing the hand associated with that symbol. We did not punish the monkeys for errors. The monkeys were trained to only release one hand in response to the presented symbol; if they released both hands, the trial was automatically aborted. We trained the monkeys to associate a specific pair of symbols (green square and pink square) with specific actions (left and right-hand release, respectively) for about 4-6 months until their performance was above 95% correct; we refer to this as the overtrained condition.

In the visuomotor associative learning version of the task (**Fig 1**), we began every recording session by presenting the monkeys with the same overtrained symbol pair (overtrained condition), and after a number of trials, switched the symbol pair to two fractal symbols, which the monkey had never seen before (novel condition), and did not have colors matching the overtrained symbols. Over ∼50-70 trials, the monkey gradually learned which symbol was associated with which hand. The manipulanda remained the same throughout the task. In the manipulanda change task (**Fig 2**), we began every recording session by presenting the monkeys with the same over-trained symbol pair and bar manipulanda, and after a number of trials, switched the bar manipulanda to dowel manipulanda. The visuomotor association remained the same throughout the task.

#### Stimulus-reward association task

The task began with the monkeys grasping two bars, one with each hand, after which a red 1° x 1° square appeared as a trial cue for 800 ms. We rewarded the monkeys with a drop of juice for not moving their hands or releasing the hand at the end of the 800 ms delay.

We did not punish the monkeys for errors and the monkeys were free to move their eyes in all the task paradigms.

### Data collection

#### Single unit recording

We introduced glass-coated tungsten electrodes with an impedance of 0.8-1.2 MOhms (FHC) into the left mid-lateral cerebellum of monkeys every day that we recorded using a Hitachi microdrive. We passed the raw electrode signal through a FHC Neurocraft head stage, and amplifier, and filtered through a Krohn-Hite filter (bandpass: lowpass 300 Hz to highpass 10 kHz Butterworth), then through a Micro 1401 system, CED electronics. We used the NEI REX-VEX system coupled with Spike2 (CED electronics) for event and neural data acquisition. We verified all recordings off-line to ensure that we had isolated Purkinje cells (**Fig 1d**) and that the spike waveforms had not changed throughout the course of each experiment. We identified cerebellar Purkinje cells online by the presence of complex spikes online, and offline by the i) spike waveforms, ii) a pause in simple spike after a complex spike, and iii) the simple spike interspike interval distribution(*38*).

#### Hand tracking

We either painted a spot on the monkeys’ right hand with a UV-blacklight reactive paint (Neon Glow Blacklight Body Paint) prior to every session or tattooed the right hand with a spot of UV Black light tattoo ink (Millennium Mom’s Nuclear UV Blacklight Tattoo Ink). We used a 5W DC converted UV black light illuminator to shine light on the spot. Then we used a high speed (250 fps) camera (Edmund Optics), mechanically fixed to the primate chair, to capture a video sequence of the hand movement while the monkeys performed the tasks. We only tracked the monkeys’ right-hand movement because the neurons had similar response with either hand movement (**Fig S1d**).

#### Licking

We recorded licking at a sampling rate of 1000 Hz using a capacitive touch sensor coupled to the metal water spout which delivered liquid water reward near the monkey’s mouth. Raw binary lick traces were used to generate instantaneous lick rate by trial averaging and smoothing it with a Gaussian kernel of sigma = 20 ms.

#### Eye movements

We tracked the monkey’s left eye positions at 240 Hz sampling rate with an infrared pupil tracker (ISCAN, Woburn, MA USA) interfaced with Spike 2 (CED electronics) where it was upsampled to 1000 Hz and synced with the event markers from NEI REX-VEX system.

#### Magnetic Resonance Imaging

We recorded the activity of a task-related P-cell, removed the microdrive, and secured the electrode to the guide tube with a dab of acrylic. We transported the animal to the MRI center in the same building as the recording lab, anesthetized him as for surgery, and held him in the 3T Siemens MRI scanner using a Kopf MRI compatible stereotaxic apparatus. We estimated the lobules from an adjacent slice which did not have the electrode artifact, and extrapolated the lobule identity.

### Data Analysis

#### Preference indexes

We defined hand preference index (HPI) and symbol preference index (SPI) as the normalized difference in spiking activity between left and right hands or the two symbols presented during the over-trained task.

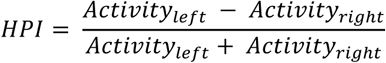

similarly,

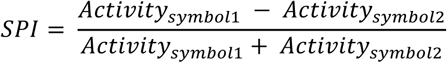

These indexes range from −1 to 1.

#### Hand movement data analysis

We used the track mate feature(*39, 40*) and custom written software in MATLAB to semi-manually track the fluorescent paint spot painted on the monkey’s hand. First, we used a downsampled LoG (Laplacian of Gaussian) filter, usually with an estimated blob diameter of 20 and a threshold of 20, to detect the fluorescent dot on the hand. Then we chose a range of threshold manually to detect the dot with considerable fidelity. We then tracked the spot using a nearest neighbor search approach. Briefly, the search algorithm relies on the KD-tree technique whereby the spots in the target frame are searched for the nearest neighbor of each spot in the source frame. If the spots found are closer than the maximal allowed distance (15 pixels), a link between the two spots is created and the process is repeated. We further confirmed the tracked path by using a linear assignment problem (LAP) tracker(*41*), allowing for gap filling. We manually removed spuriously detected spots from each frame in the tracks, during post processing. We then analyzed the tracks in MATLAB using custom written software. We smoothed the raw trajectories by using low pass moving filter with filter coefficients equal to the reciprocal of the span. We aligned all trajectories to the first instance of hand release from the bars. We excluded hand trajectory outliers from our data set if we could not reliably trace the trajectories.

#### MRD method to detect changes in activity pattern

The change in activity for different P-cells occurred at different times in a trial. Given this heterogeneity in its timing, there is no way to predict the time or the magnitude of the activity change, a priori, for a randomly selected P-cell. Therefore, we wanted to use a method that is blind to the time of change, its distribution or the intrinsic properties of the neuron and more importantly, makes no a priori predictions about their properties, in such a way that, given the input (for example, activity in OT and activity in N), it gives us a logical decision (classifies) as to whether the activities were different or similar between the conditions. Since the change in activity could occur at any time of the trial for a given neuron, we used the whole trial activity (from before symbol onset through after reward) to calculate MRD. However, to avoid the trivial possibility where the two activities are different merely due to changes in reaction time latencies, we limited our analyses window to: −400 to 200 ms aligned to symbol onset, and −200 to 600 ms aligned to movement onset for each neuron. The activity aligned to symbol onset and movement onset of the k-th trial for each condition (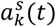 and (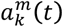 respectively) were concatenated to form one single activity vector as follows:

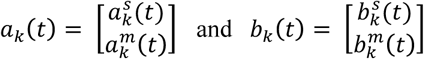

To quantify the change in activity pattern between two conditions A and B, we first identified the condition with least intra-condition variance (suppose condition A). We then compared the activity within the condition with the least intra-condition variance (A) with the activity across both conditions (A and B). Both A and B are 20 x n matrices with n being the length of the signal. First, we randomly sampled 10 trials each from the last 20 trials in condition A and the first 20 trials in the condition B and calculated the root mean squared (rms) distance between the mean activities:

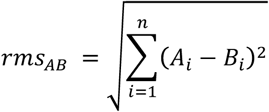

Where, 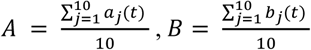 and n is the length of the signal

Also, *a*_*j*_(t) and *b*_*j*_ t were sets of 10 trials chosen randomly from 20 trials.

We repeated this process 250 times to obtain a distribution of rms distances that compared the extent of change in across-condition activity profile between conditions A and B. To compare this distribution with the within-condition activity profile, we randomly sampled 10 trials twice without replacement, from condition A (condition with the least intra-condition variance) and repeated the same analysis to obtain another distribution of rms distances.

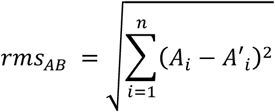

Where 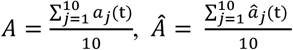 and n is the length of the signal

Also, *a*_*j*_(t) and 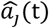 were set of 10 trials chosen randomly from N and N-10 trials respectively, where N is the number of trials in condition A.

Next, we repeated this process 250 times to get a distribution of *rms*_*AA*_ and *rms*_*AB*_. Finally, we compared means of both distributions (MRD) for statistical difference. See **Fig S2** for validity of this method applied to Gaussian distributions and Poisson spike trains. We used this method to compare both neural spike density functions and movement trajectories between different conditions.

#### Learning behavior and the criterion for ‘learned’

We constructed the learning curve for every session by calculating the percent correct trials in a sliding window of 10% of the total number of trials in that session as the bin width moved by 5% of the total number of trials in that session. If the monkeys reached >90% correct through the above method and remained above 80% for at least the next 20 trials, the associations were considered ‘learned’. All sessions where the monkeys did not reach this criterion were not used to analyze learning related trial by trial changes in activity but were retained for other analyses which did not require to study the activity through learning. Some of the earlier recorded sessions often did not start with an overtrained condition and neither did the monkeys reach the criterion for ‘learned’ and were thus excluded for studies involving changes from overtrained to novel learning and learning related changes. If the monkeys satisfactorily reached the criterion while the neuron was still stable, we proceeded to the reverse paradigm.

#### Analysis of delta epoch

##### Identification of delta epoch

We performed a t-test between pairs of time points between the mean activity traces of correct and wrong trials (sigma = 30 ms) from soon after the reward delivery (or its absence) until the next trial’s reward delivery (or its absence). We defined the start of delta epoch as the first time-point where the P value decreased below 0.05 and stayed below 0.01 continuously for the next 150 ms. We defined the end of the delta epoch as the first time-point when the P value went above 0.05 and stayed above 0.05 continuously for at least the next 250 ms.

##### Classifying P-cells into wP-cells and cP-cells

To classify the cells into wP-cells and cP-cells in an unbiased way, we took the average neural activity of correct trials and wrong trials in the delta epoch for each neuron and used different unsupervised learning algorithms such as hierarchical clustering and k-means clustering to cluster our data into two natural clusters that maximized the correlation between the data points. Hierarchical clustering finds the similarity or dissimilarity between every pair of data points and groups them into a binary, hierarchical cluster tree. Then it determines where to cut the hierarchical tree into smaller clusters. `k-means` partitions the data into *k* mutually exclusive clusters. `By` treating each data point as having an unique location in space, it finds a partition in which the distance between the data points within a given cluster is minimized while the distance between the data points in other clusters are maximized. Unlike hierarchical clustering, *k*-means clustering operates on real data, rather than on the larger set of dissimilarity measures.

We used correlational distance metric for these algorithms which is defined below: Let c be the centroid of a cluster and x be individual data points. Then we define the correlational distance d(x,c) as:

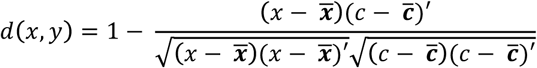

Where, 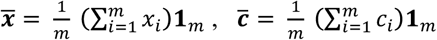 and 1_*m*_ is a 1 x m unit vector.

We obtained very similar results from both these disparate clustering algorithms. More importantly, both types of clustering lead to very similar results as simply classifying the cells based on the trial type with maximum average neural activity despite the difference in neural activity between the two trial types (92% similarity for k-means and 100% similarity for hierarchical clustering). We labelled the P-cells as wP-cells and cP-cells based on these results. Thus, this provided an unbiased way of classifying cells.

##### Effect of N-2 and N+1 decisions

Having labelled the P-cells, for each neuron, we calculated the average neural activity in the two-back decision (2-N) case and the impending decision (1+N) case, without changing the labels. For both these cases, the clusters (corresponding to wP-cells and cP-cells obtained previously) were intermixed without any distinction and the same algorithms failed to partition the P-cells into clusters in both these cases (**Fig 4F-H**).

#### Statistics

To check if two independent distributions were significantly different from each other, we first performed a two-sided goodness of fit Lilliefors test, to test for normality, then used an appropriate t-test; or else a non-parametric Wilcoxon ranksum test. All error bars and shading in this study, unless stated otherwise, are mean ± s.e.m.

#### Reinforcement-drift diffusion model

Consider learning one symbol-action-outcome association for one symbol with two alternative action choices, *a*_*L*_ and *a*_*R*_ corresponding to left and right hand bar-release hand movements respectively. We model the action selection through a race to threshold model where there is a race in the evidence accumulation between the action values *a*_*L*_ and *a*_*R*_ modeled by Wiener first-passage time (WFPT) distribution. This calculates the likelihood of the reaction time of choice i given by:

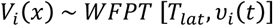

where *T*_*lat*_ is a non-decision time parameter that captures the latency of stimulus encoding and/or motor processes. We assume it to be constant throughout learning and fixed for either action value and *v(t)* denotes the rate of accumulation process for the trial *t*. Therefore, we have two independent action choice accumulators for left and right action choices.

The rate of accumulation in the overtrained (OT) condition, 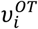 is assumed to be a constant that does not change with trial. Then, we model the evolution of *v*_*i*_ *t*, with learning as:

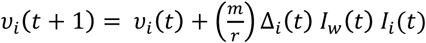

Where

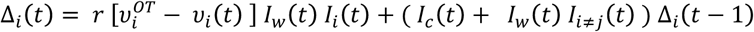

here, *m* captures the rate of learning or the proportion of the error that is compensated for from trial to the next trial. The value of *m* tends to be between 0.10 and 0.30, meaning trial-to-trial corrections adjust for approximately 10–30 % of the error. *r* is the scale factor, *I*_*i*_ (*t*) is the indicator function describing the chosen action (*a*_*ch*_) on trial *t* defined as follows:

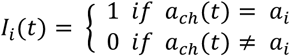

*I*_*w*_(*t*) and *I*_*c*_(*t*) are indicator functions describing the outcome of trial *t*, defined as follows:

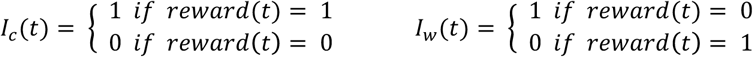

We propose that the rate of change of the accumulation rate *v*_*i*_(*t*), given by Δ_*i*_(*t*) is represented in the P-cell activity of the delta epoch. Clearly, the delta activity is updated as the scaled difference between the rate of accumulator in the OT condition and that in a given trial, if the trial were wrong and the monkey chose the hand corresponding to the accumulator. Else, the delta value remains the same.

Similar to previous observation, the monkeys learned the two stimulus-action-reward associations roughly independently(*42*). Therefore, we modeled acquisition of each stimulus-action-reward association independently with negligible interference between the associations. Furthermore, we only model the magnitude of the activity in delta epoch for each P-cell and its evolution with learning because we see similar changes in both cP-cells and wP-cells from our analyses.

Since we only had ∼30 trials in the OT condition and roughly 50-70 trials for the entire learning condition per session, we pooled data across those sessions in which the reaction time increased significantly, to increase statistical power and get enough data points to fit the data optimally. Therefore, our model was only fit to the population data and not individual sessions. First, we uniformly and coarsely sampled a range of values that could generate the observed RT distributions, to simulate RT distributions of 2,000 trials, using the following equation:

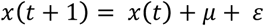

where *x*(*t*) is the level of accumulation at time *t*; *μ* is the mean drift rate and *ε* is a Gaussian noise term with mean = 0 and standard deviation that represents the noise in the input signal. We modeled the process to be ideal, with a time step of 1 ms, without any leak term. The stochastic nature of the process generates a RT distribution, that changes with *μ*. The variability of the accumulation process at a given time *t* is given by:

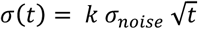

where *σ* is the standard deviation of the process, *k* is a constant scaling factor and *σ*_*noise*_ is Gaussian noise.

Second, we used MATLAB’s fmincon function for minimize the error between the generated and the empirical RT distributions(*43*). We applied a nonlinear constraint which restricted at least 70% of the simulated RT distribution to lie within the extent of the empirical RT distribution. We then calculated the error as the difference between inverted Gaussian weighted CDF of the empirical and simulated RT distributions. This enabled better fits to the tails of the RT distributions to obtain a more reliable estimate of the standard deviation of the distributions. Although the minimum error solution typically converged within <30 iterations for *μ* and *σ*, we ran the minimization process multiple times with different starting points to ensure that we obtained the best parameters. We performed this process for two sections of learning: in the OT and initial learning for left and right hand RTs. This gave us estimates of 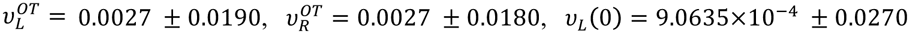 and *v*_*R*_ (0) = 9.8756×10^−4^ ± 0.0250. We then simulated the learning process with these values and compared the simulated learning curve and the neural activity in the delta epoch to the experimental observation (**Fig S11**).

**Figure S1:**
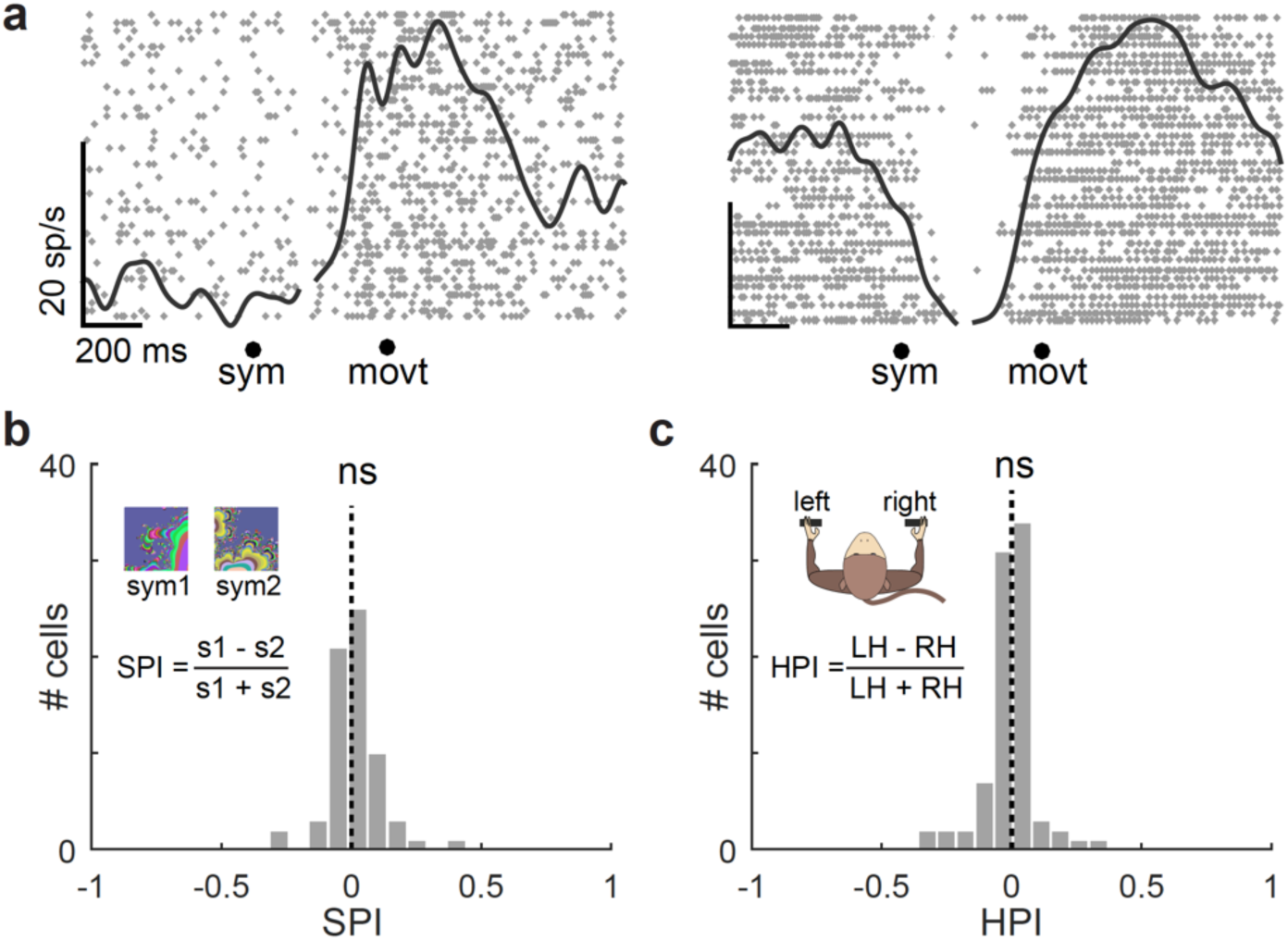
P-cells showed increased activity during the overtrained task. **a**. Two representative P-cells that showed an elevated activity during hand movement. The cell on right, in addition, showed a stimulus dependent decrease in activity. **b**. Histogram of Symbol Preference Index (SPI) calculated for all neurons. P = 0.3326; n.s. means not significantly different from 0; t-test. **c**. Same as h, but for hand preference index (HPI). P = 0.5787; t-test; n.s. means not significantly different from 0; t-test.

**Figure S2:**
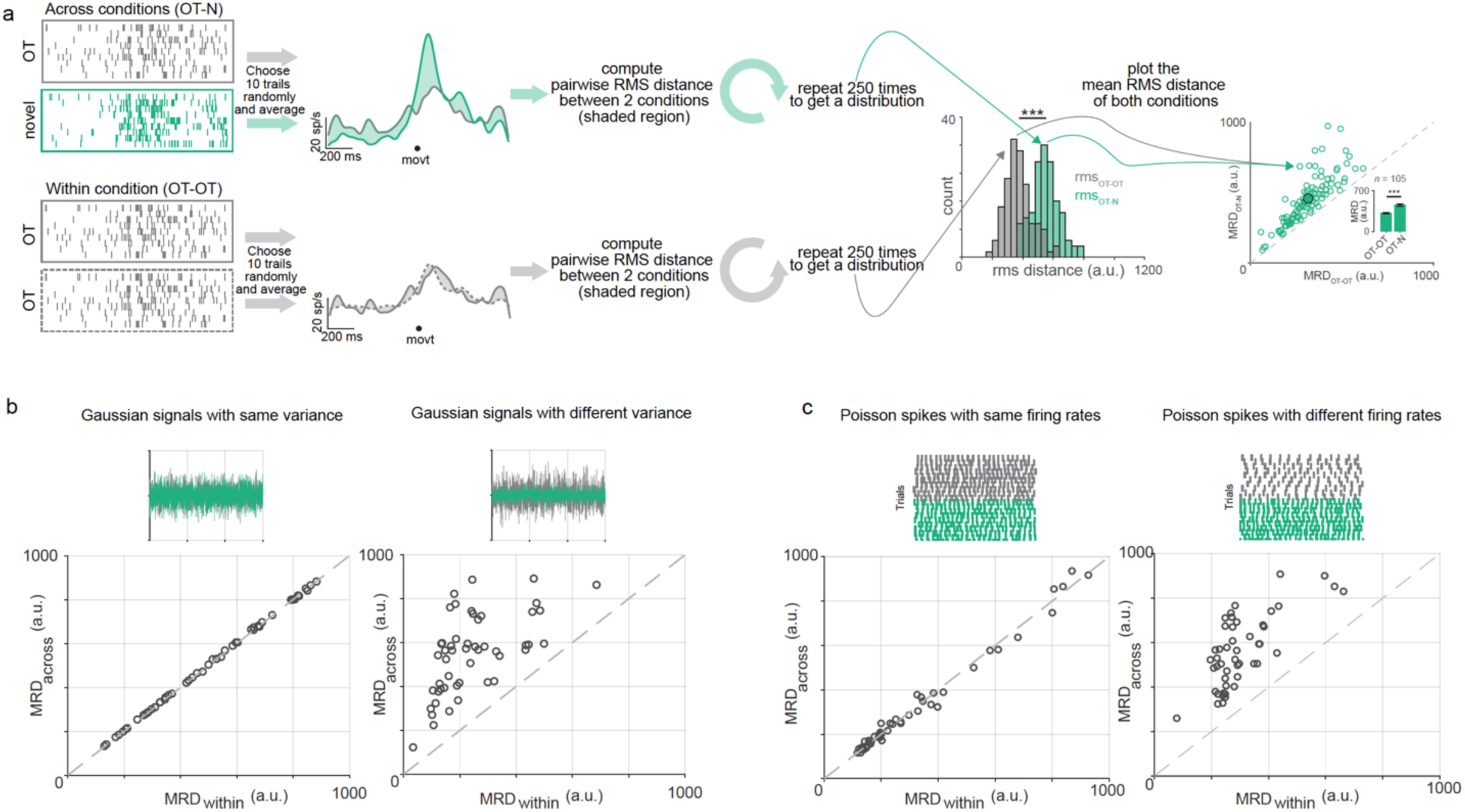
MRD method. **a. Top:** First, we randomly sampled 10 trials each from the last 20 trials in the overtrained condition (grey) and the first 20 trials in the novel condition (green) and calculated the root mean squared (rms) distance between the mean activities. We repeated this process 250 times to obtain a distribution of rms distances that compared the extent of change in across-condition activity profile in the novel condition from the activity profile in the overtrained condition. **Bottom:** To compare this distribution with a control null-distribution, we randomly sampled 10 trials twice without replacement from the overtrained condition and repeated the same analysis to obtain another distribution of rms distances to obtain an estimate of variability of within-condition. A simple test of statistical significance between the mean of these two distributions would provide an estimate of the change between conditions. **b**. Left: MRD Method applied to 50 pairs of Gaussian signals with 0 mean and same variance. Top panel shows one such pair. Bottom panel shows the Mean RMS Distance (MRD) calculated across group plotted against that calculated within the first group. Right: Same as before, but for 50 pairs of Gaussian signals with 0 mean and different variances. **c**. Left: Same as b, but for 50 pairs of mean spike density functions obtained from Poisson spike trains with same mean and variance (λ). Same as b, but for 50 pairs of mean spike density functions obtained from Poisson spike trains with different means and variances (λ).

**Figure S3:**
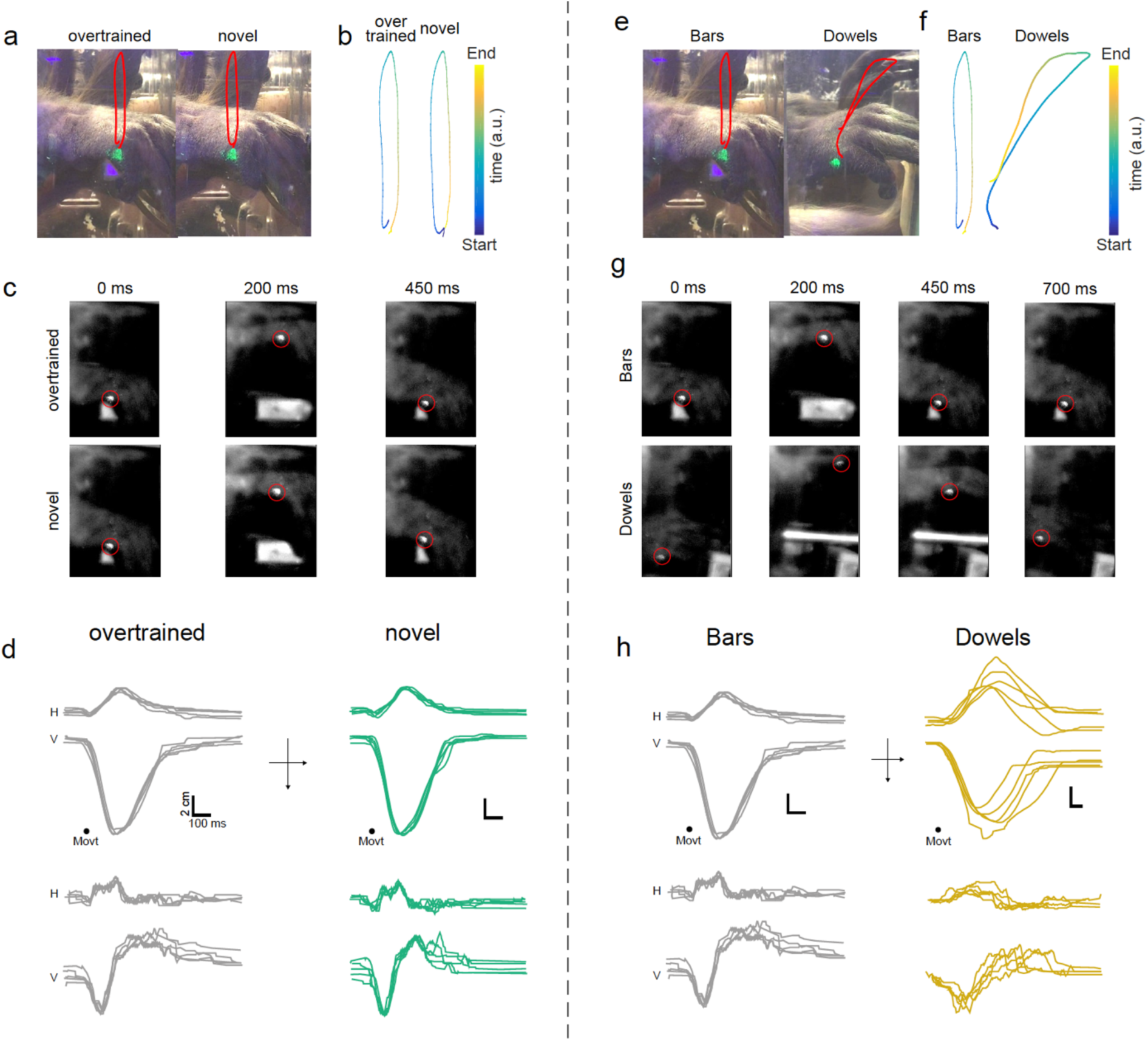
Motor kinematics. **a**. Schematic of the hand movement trajectory as the monkey lifted its hand off the bar in the overtrained (left) and novel conditions (right). The red trace maps the position of the green fluorescent marker in space through the hand movement. **b**. Position of the fluorescent marker in space and time (color indicates the time as in the colorbar inset). **c.** Snapshots from high-frame rate movies showing the monkey’s hand movement trajectory at three time points (0, 400 and 900 ms from the start of movement) for overtrained (top three panels) and novel (bottom three panels) conditions. Red circle marks the fluorescent marker. **d**. Top panel: Horizontal (H) and Vertical (V) hand trajectories for five continuous trials aligned on movement onset for overtrained condition. Bottom panel shows the H and V velocities for the same five trials. **e**. Schematic of the hand movement trajectory during bar lift (left) and the dowel release (right) conditions. The red trace maps the position of the green fluorescent marker in space through the hand movement. **f**. Position of the fluorescent marker in space and time (color indicates the time as in the color bar inset). **g**. Snapshots from high-frame rate movies showing the monkey’s hand movement trajectory at four time points (0, 400 and 900 and 1400 ms from the start of movement) for bar lift (top three panels) and dowel release (bottom three panels) conditions. Red circle marks the fluorescent marker. **h**. Top panel: Horizontal (H) and Vertical (V) hand trajectories for five continuous trials with, bar release, aligned on movement onset. Bottom panel shows the H and V velocities for the same five trials.

**Figure S4:**
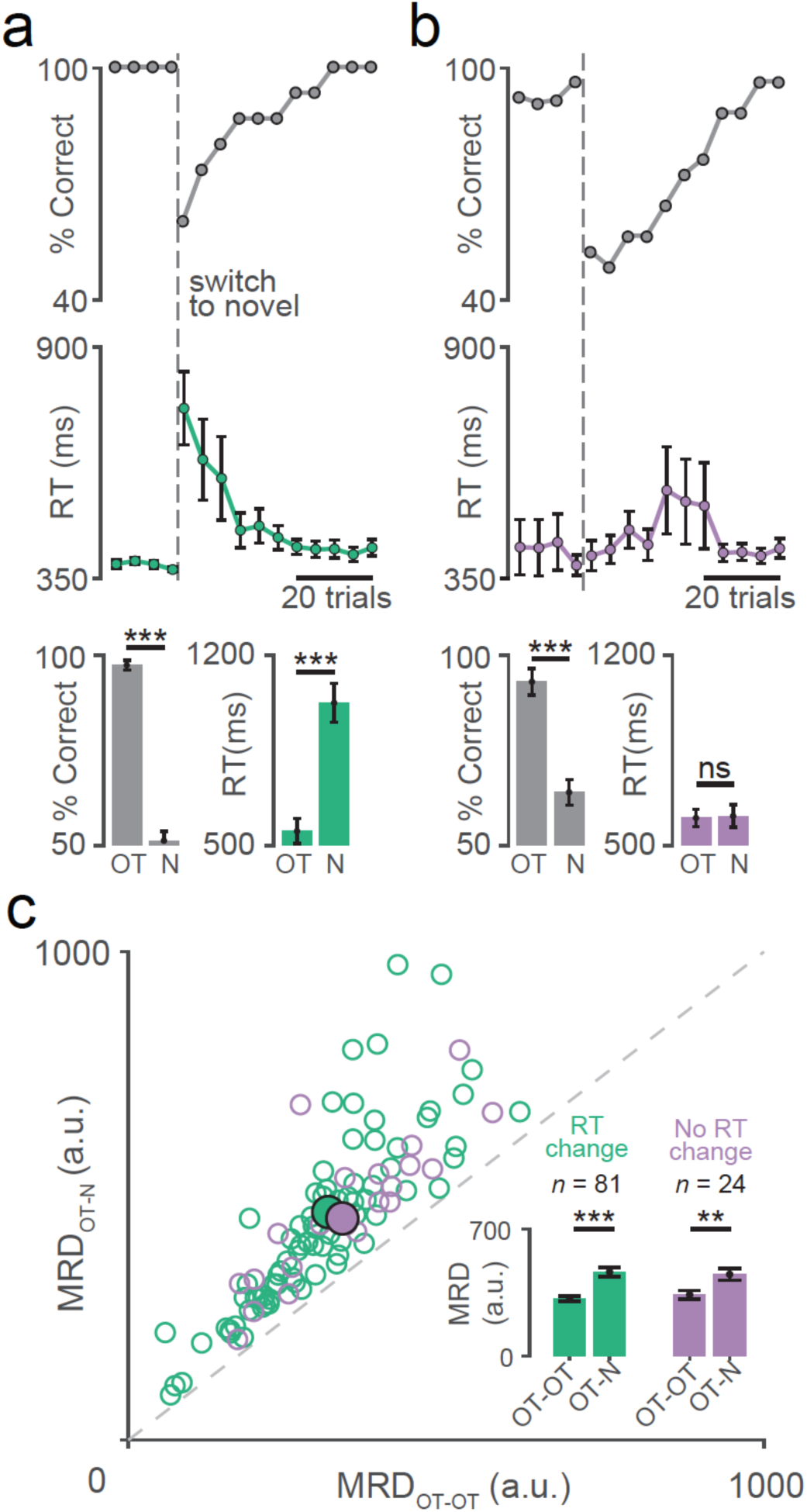
Neural activity changes were independent of changes in reaction time. **a**. Top: Percent of correct trials plotted as a function of trial number relative to the switch to novel visuomotor association. Middle: Reaction times for the same trials. Error bars show the standard error of the mean. Bottom: Mean percent correct (left) and reaction time (right) in the overtrained (OT) and novel (N) conditions for all the sessions with changes in reaction time (*** means *P* = 10^−14^, Wilcoxon rank sum test). **b**. Percent correct(top), reaction time (middle), and session averages(bottom) for sessions in which the manual reaction time did not change (n.s means *P* = 0.8151, Wilcoxon rank sum test) after the switch to novel visuomotor association but the performance did (*** means *P* = 10^−7^, Mann-Whitney U-test). Same convention as a. **c**. Same data as in Fig 1c but separated into sessions with RT change (green; *** means *P* = 10^−6^, Mann-Whitney U-test) and no RT change (violet; ** means *P* = 10^−3^, t-test).

**Figure S5:**
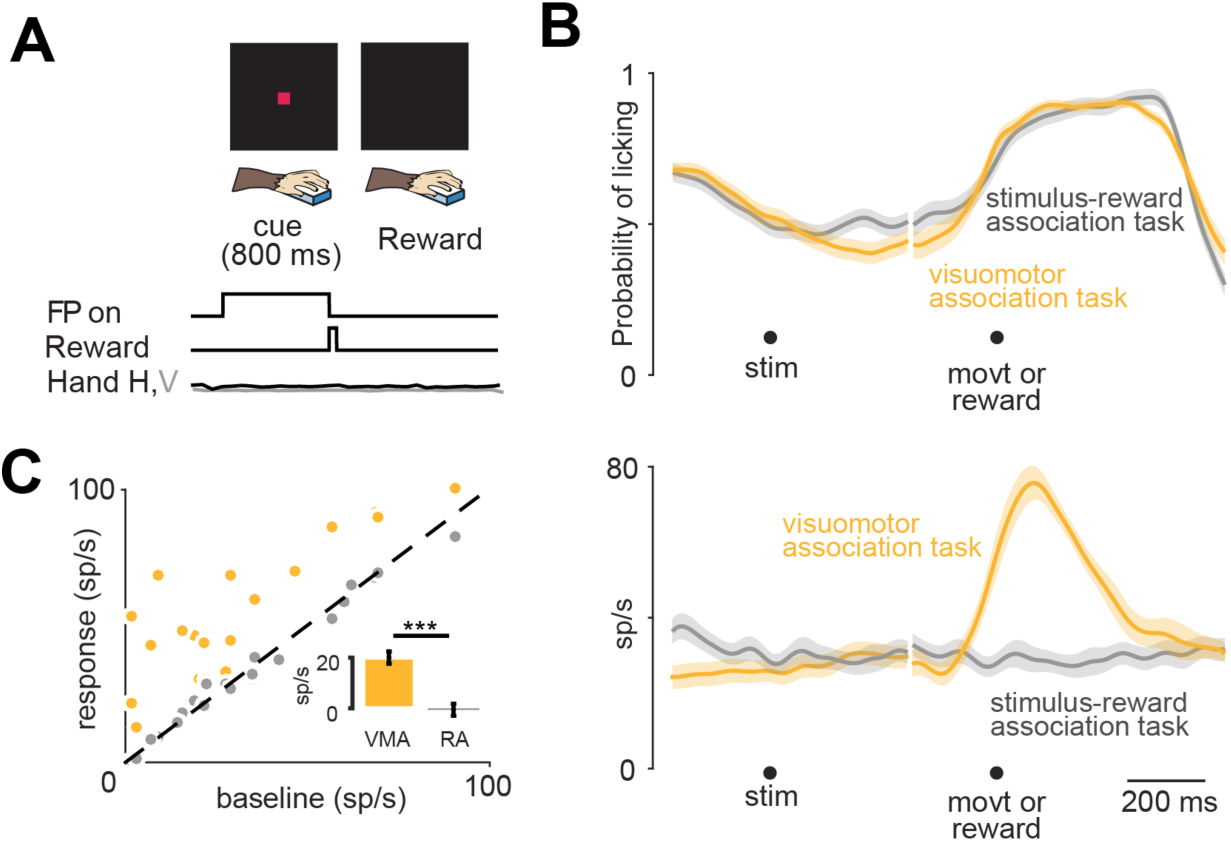
P-cells did not respond in a stimulus-reward association task. A. Stimulus-reward association task: A red square (cue for the start of the trial) appeared in the center of the screen for a fixed duration of 800 ms and then disappeared as the monkeys received a liquid reward if they made no hand movement. B. Top: The monkeys started licking the juice spout in the same way during the reward anticipation task (grey) as well as the visuomotor association task (gold). Bottom: A representative P-cell showing a movement related increase in activity during the hand movement in the visuomotor association task while showing no significant modulation in activity during a stimulus-reward association task. C. aBaseline activity plotted against the peak firing rate in the movement epoch during the visuomotor association task (gold) and the stimulus-reward association task (grey) for all 25 P-cell. Inset shows the mean difference in peak activity from baseline firing for the two conditions. *** means P = 10^−7^, Mann-Whitney U-test.

**Figure S6:**
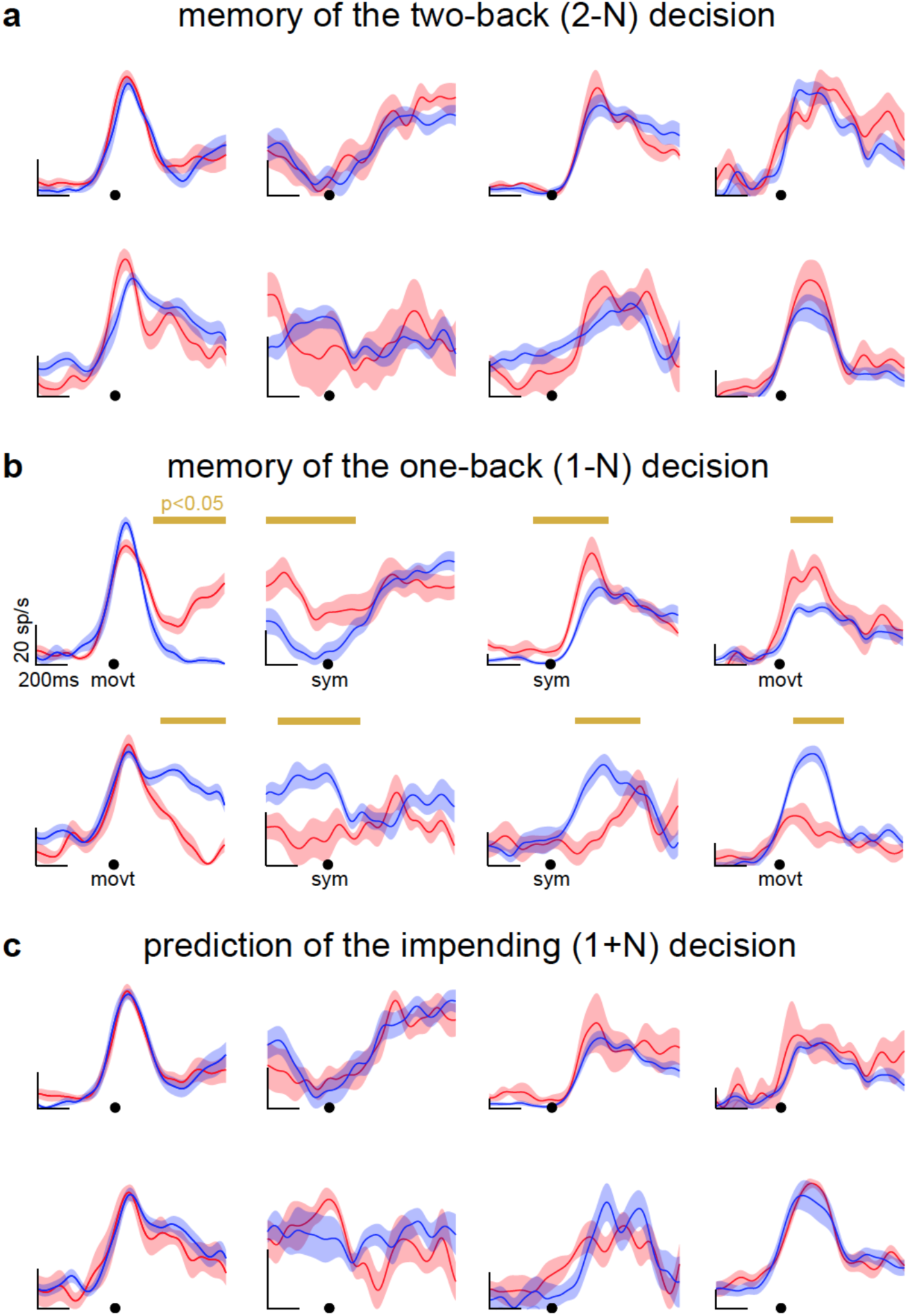
Delta epoch only encoded the memory of the most recent decision. Same P-cells as in Fig 4a-b analyzed on activity of two back decision (2-N) (a), one back decision (1-N) (b) and impending decision (N + 1)(c). Note that panel b is same as **Fig 4 A-B**.

**Figure S7:**
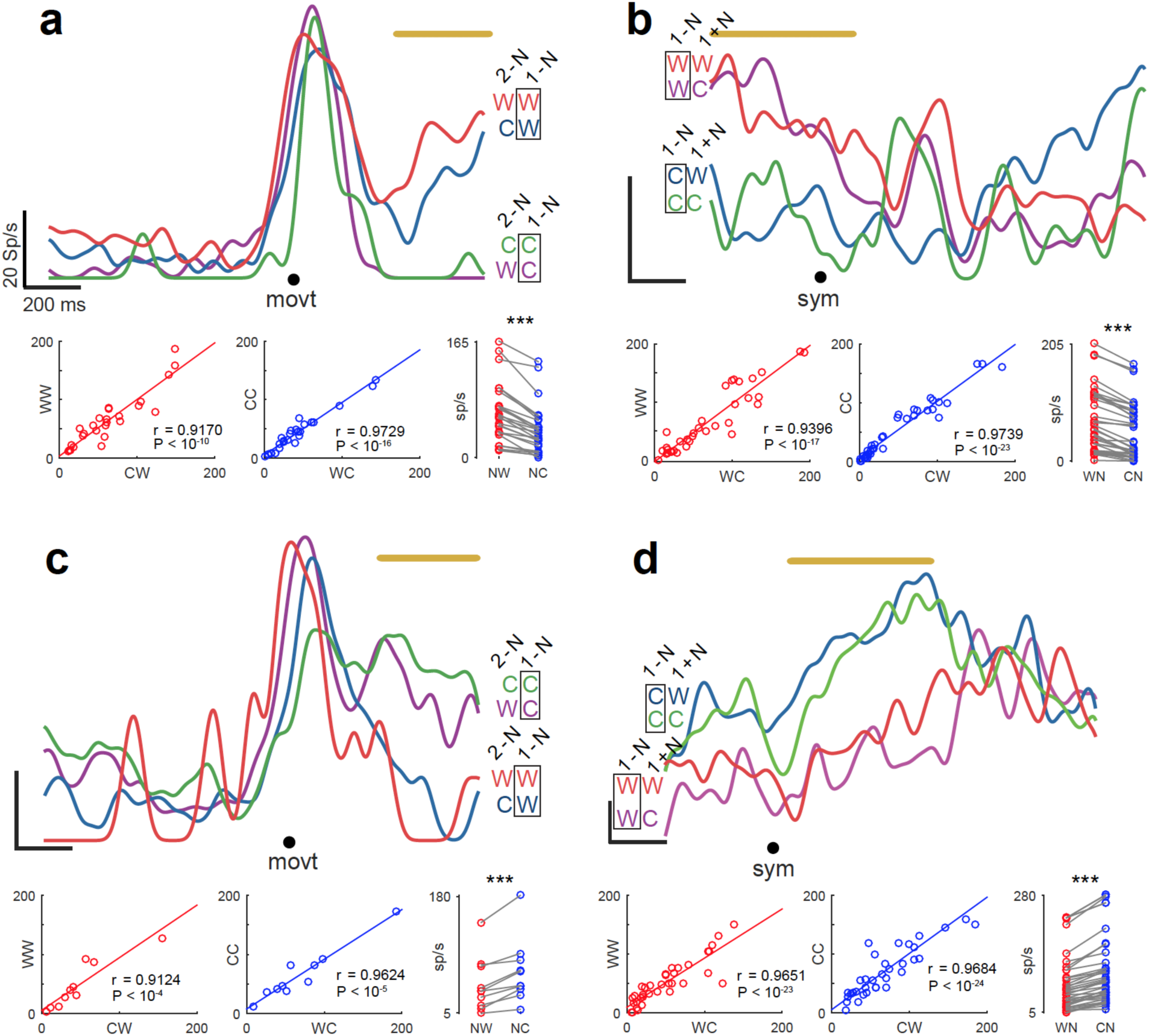
Memory of a single decision. a. Top: Movement aligned spike density function of wP-cell from **Fig 4a leftmost panel**, separated into two-back (2-N) and one-back (1-N) decisions. Black rectangle highlights the fact that this P-cell had similar activity for 1-N decision regardless of the 2-N decision. Neural activity in delta epoch for CW condition was significantly correlated with WW condition (ho = 0.9170; P<10^-10^) and was not significantly different (P = 0.8447; Mann Whitney U test). Bottom center: Neural activity in delta epoch for WC condition was significantly correlated with CC condition (rho = 0.9729; P<10^-16^) and was not significantly different (P = 0.6575; Mann Whitney U test). Bottom right: Activity in one-back wrong trials (NW) was significantly higher than activity in one-back correct trials (NC). Taken together, the neural activity was similar for 1-N decisions regardless of the 2-N decisions. In other words, P-cells did not have a history of the outcome of prior decisions other than the 1-N decision. b. Top: Same as a, but for the w-Pcell from **Fig 4a left center panel**, separated into one-back (1-N) and no-back (0-N) decisions. Black rectangle highlights the fact that this P-cell had similar activity for 1-N decision regardless of the 1+N decision. Bottom left: Neural activity in delta epoch for WC condition was significantly correlated with WW condition (rho = 0.9396; P<10^-17^) and the mean was not significantly different (P = 0.6276; Mann Whitney U test). Bottom center: Neural activity in delta epoch for the CW condition was significantly correlated with CC condition (rho = 0.9739; P<10^-23^) and was not significantly different (P = 0.5737; Mann Whitney U test). Each marker is a neuron. Bottom right: Activity in one-back wrong trials (WN) was significantly higher than activity in one-back correct trials (CN). *** means P<0.001, t-test. Taken together, the neural activity was similar for 1-N decisions regardless of the 0-N decisions. In other words, P-cells could not predict the outcome of impending decisions. c. Top: Same as a, but for the cP-cell from **Fig 4b leftmost panel**. Bottom left: Neural activity in delta epoch for CW condition was significantly correlated with WW condition (rho = 0.9124; P<10^-4^) and was not significantly different (P = 0.9965; Mann Whitney U test). Bottom center: Neural activity in delta epoch for WC condition was significantly correlated (rho = 0.9624; P<10^-5^) with CC condition and was not significantly different (P = 0.8771; Mann Whitney U test). Bottom right: Activity in one-back wrong trials (NW) was significantly lower than activity in one-back correct trials (NC). d. Top: Same as b, but for the cP-cell from **Fig 4b right center panel**. Bottom left: Neural activity in delta epoch for WC condition was significantly correlated with WW condition (rho = 0.9651; P<10^-23^) and was not significantly different (P = 0.5347; Mann Whitney U test). Bottom center: Neural activity in delta epoch for CW condition was significantly correlated with CC condition (rho = 0.9684; P<10^-24^) and was not significantly different (P = 0.8920; Mann Whitney U test). Bottom right: Activity in one-back wrong trials (WN) was significantly lower than activity in one-back correct trials (CN).

**Figure S8:**
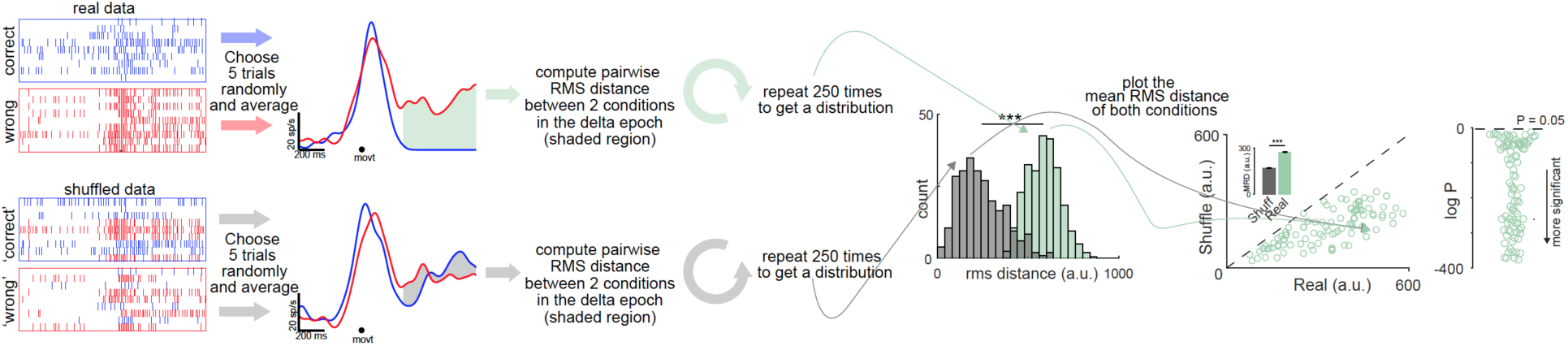
Delta epoch did not arise by chance. First, we randomly sampled 5 correct trials and 5 wrong trials from the first 20 trials in the learning condition and calculated the root mean squared (rms) distance between the mean activities in the delta epoch (top row to the left). We repeated this process 250 times to obtain a distribution of rms distances that provided an estimate of the true difference between the neural activity due to correct and wrong trials in the delta epoch (green histogram in the middle). We then shuffled the label of ‘correct’ and ‘wrong trials’ amongst the first 20 learning trials (bottom row to the left) and repeated the same procedure as above, 250 times, on the new correct and wrong trials, created during each iteration. This provided the null estimate (grey histogram in the middle). A simple test of statistical significance between the mean of these two distributions (scatter plot to the right) would tell us if the delta epoch was a true phenomenon or happened due to chance. The rightmost plot shows the p value from t-test of each comparison in the scatter plot.

**Figure S7:**
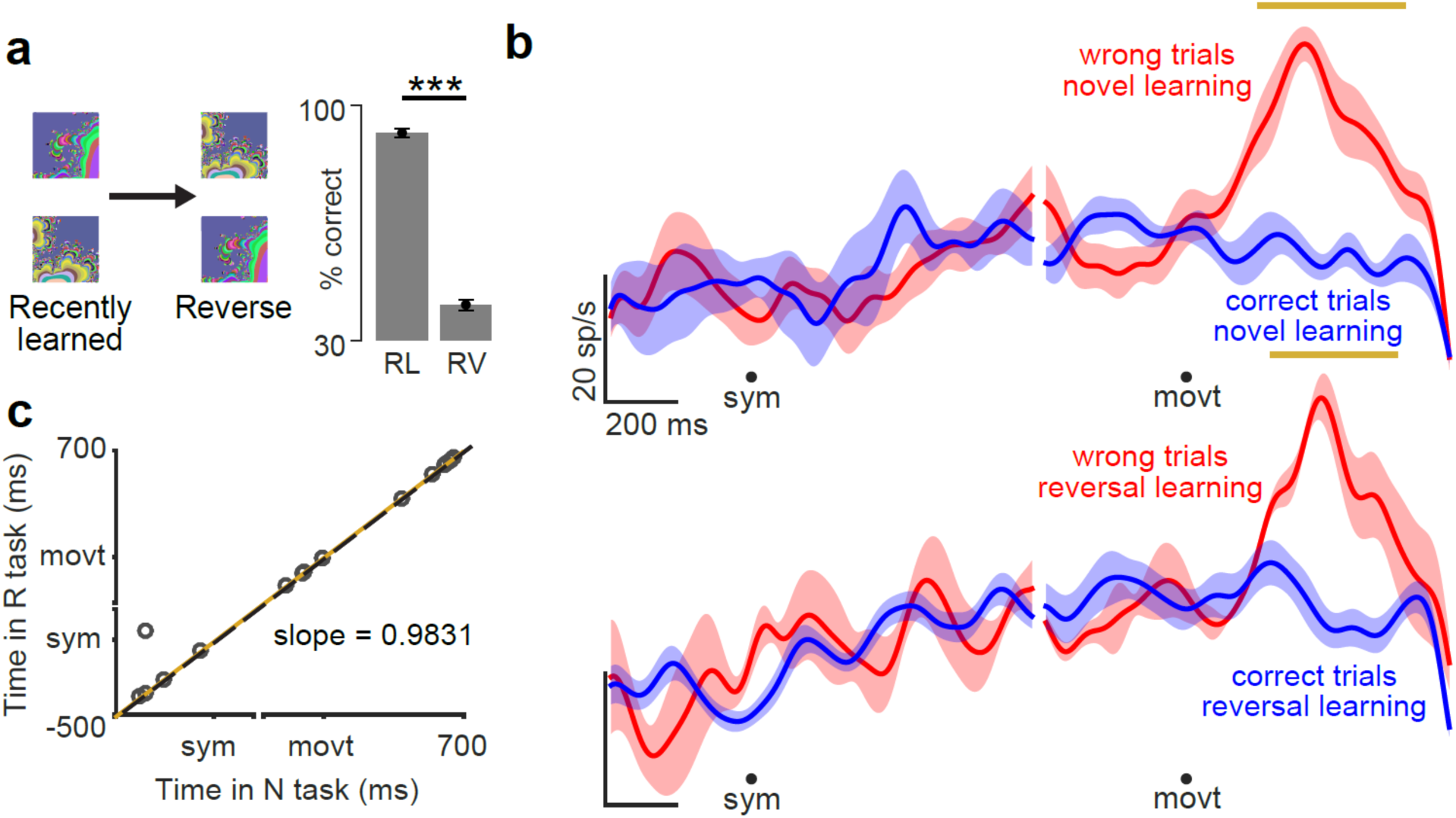
Timing of delta epoch is consistent in novel and reversal learning. a. Task in which the symbol-hand associations were reversed after the monkeys learned the novel associations. Right: Average learning rate for last few recently learned (RL) novel associations trials and first few reversal (RV) trials *** means P = 10^−7^, Mann-Whitney U-test. b. A representative P-cells showing delta epoch (indicated by gold line) in the same interval during novel learning (top) and reversal learning (bottom). c. The mean time of delta epoch for each cell (each marker) in the novel learning (abscissa) vs reversal learning (ordinate) conditions. Broken line is the line of unity. Solid gold line is the least squared fit line with least square slope = 0.9831 rho = 0.9709 and P=10^-10^. *N*=24 neurons.

**Figure S10:**
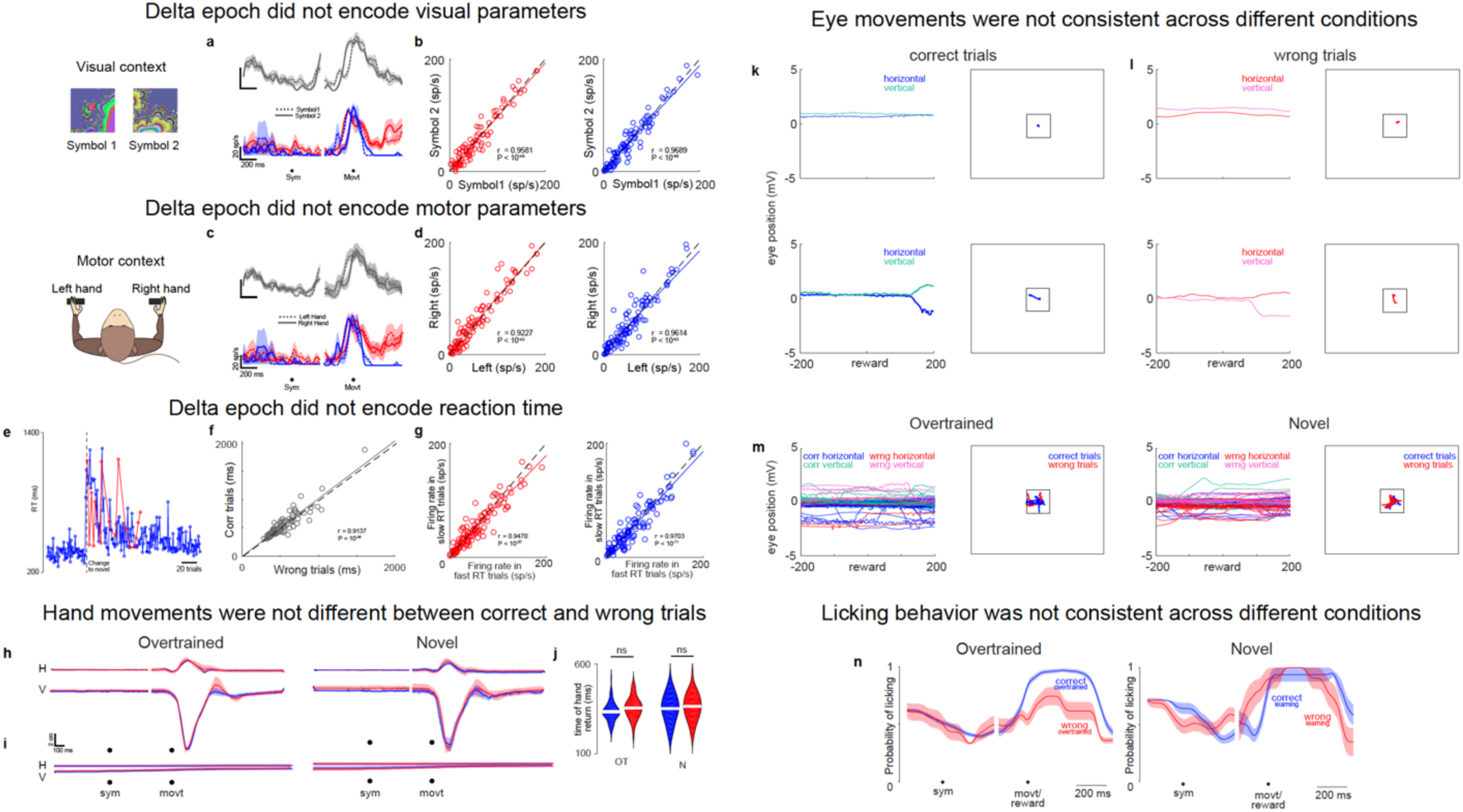
Evidence for independence of neural activity in the delta epoch from sensorimotor parameters of the task. **a**. Top: Spike density functions of a representative neuron’s activity in the overtrained condition for symbol1 (dotted line) and symbol2 (solid line) aligned to symbol onset and movement onset. Bottom: Same neuron as top left panel; but neural activity during learning for symbol1 and symbol2, separated into correct trials (blue) and wrong trial (red). **b**. Left panel: Neural activity in delta epoch of wrong trials during learning for symbol1 (abscissa) vs symbol2 (ordinate). Dashed line is the line of unity. Solid red line is the least squared fit line. Right Panel: same as left panel, for neural activity in delta epochs of correct trials. This indicates that the neural activity during learning has no symbol preference in correct (P = 0.7090; Mann Whitney U test) and/or wrong trials (P = 0.8178; Mann Whitney U test). **c**. Same as a, but for hand preference index (HPI). P = 0.5787; t-test. This means the neural activity in the overtrained condition had no hand preference. **d**. Same as b, for left and right hands, indicating that the neural activity during learning had no hand preference in correct and/or wrong trials. **e**. Reaction time for all the trials (correct: blue; wrong: red) for one representative session showing that the reaction time for correct and wrong trials during initial learning were comparable and showed a similar trend **f**. Reaction time for wrong trials (abscissa) plotted against the reaction time for correct trials (ordinate). Each dot is a session. Dashed line is the line of unity and solid grey line is the least squared fit. This means that the reaction for correct and wrong trials during initial learning were statistically comparable (P = 0.8000; Mann Whitney U test). **g**. Left: Firing rate for all cells in the delta epoch for wrong trials in the fast reaction time trials (abscissa) vs slow reaction time trials (ordinate). Each dot is a cell. Dashed line is the line of unity and solid grey line is the least squared fit. This means that the neural activity in wrong trials was not significantly different between fast and slow reaction time conditions (P = 0.7772; Mann Whitney U test). Right: Same as left but for neural activity in correct trials (P = 0.6306; Mann Whitney U test). **h**. The hand movements after correct and wrong trials were similar for the hand that was used to report the choice in the overtrained (left) and novel (right) conditions. **i** The hand movements after correct and wrong trials were similar for the hand that was not used to report the choice in the overtrained (left) and novel (right) conditions. Note that ‘movt’ here represents the time at which the other hand movement was initiated. **j**. The time of hand return to the bar was similar after correct and wrong trials during overtrained (P = 0.3663, ranksum test) and novel (P = 0.1452, ranksum test) conditions. **k**. Mostly, the monkeys fixated at the center (top) while sometimes they made small spontaneous eye movements (bottom). Eye movements shown here are for correct trials. Left panel shows the horizontal and vertical components of the eye positions aligned on the reward onset. Right panel shows the actual eye movement constructed from the horizontal and vertical components from the left panel. We always flashed the stimulus (cue or the symbols) only at the center of the screen and the monkeys were free to move their eyes. Black square represents the symbol. **l**. Same as a; but for eye movements for wrong trials. **m**. Since the monkey’s eye movements were not constrained in any way, they made reward independent-free eye movements and therefore, their eye movements did not have a consistent pattern for correct (blue) or wrong trials (red) in OT (left) or N condition (right). However, the neural activity significantly differed between correct and wrong trials in the delta epoch. Taken together, this suggests that the neural activity in the delta epoch was independent of eye movements. **n** In the OT task, the monkeys occasionally made errors. The monkeys licked for correct trials more than for the wrong trials (left). However, the neural activity was the same for correct and wrong trials (**Fig 3e-f**). Moreover, in the novel condition, during early learning, since the monkeys are merely guessing, the licked after they had made a hand movement ‘expecting’ a reward regardless of the task outcome and hence their licking behavior looked similar for correct and wrong trials (right). Despite this similarity in licking, we see the neural activity differ between correct and wrong trials in the delta epoch (**Fig 3a-d**). Taken together, these results strongly suggest that the neural activity in the delta epoch was independent of changes in licking behavior.

**Figure S11:**
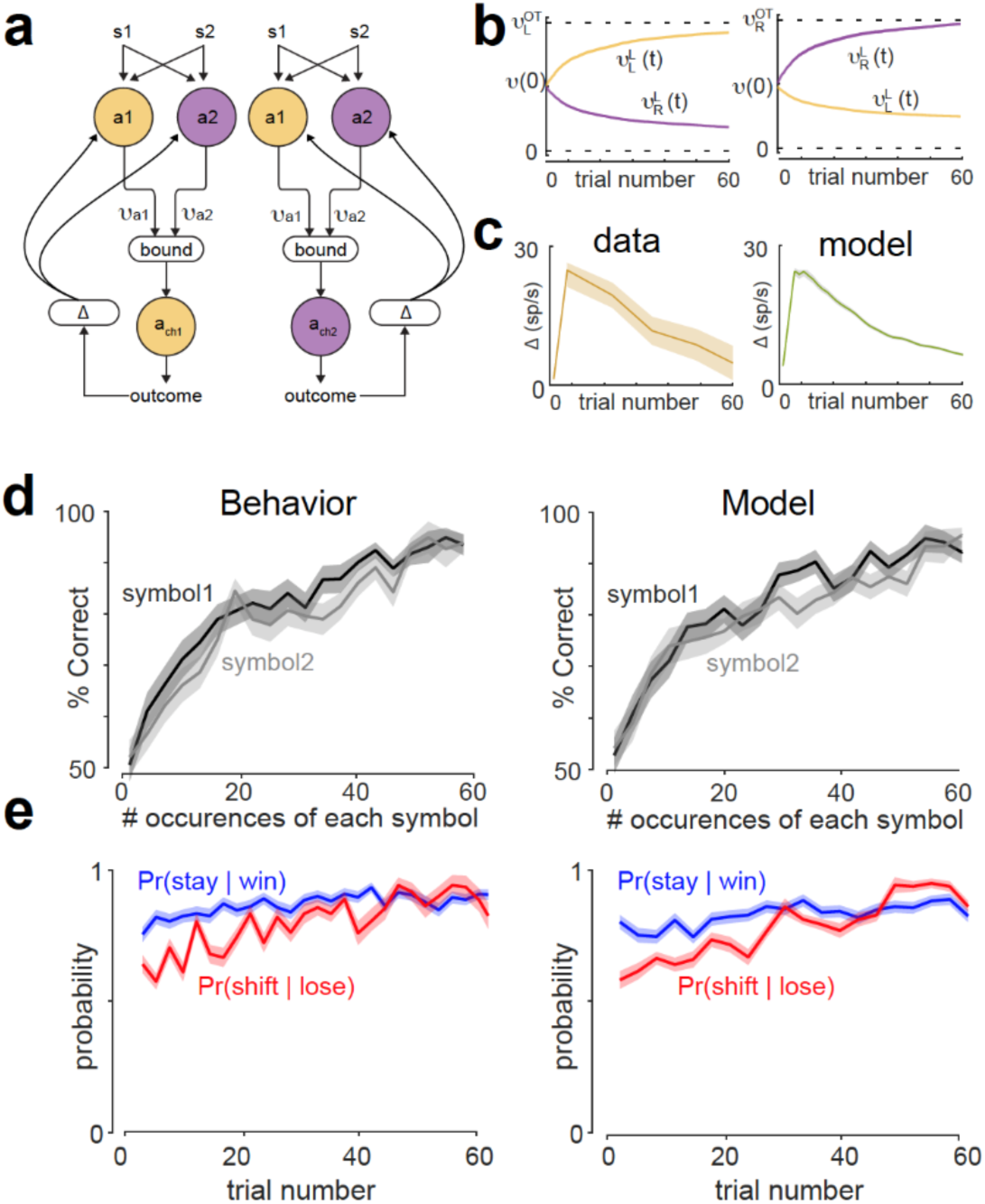
Reinforcement-drift diffusion model. a. Schematic of the model, a1 and a2 are the two action choices that are modelled as accumulators with rates *v*_a1_ and *v*_a2_ respectively, racing to threshold (bound). The winner takes all and consequences of the chosen action a_ch_ is evaluated by the activity of P-cells in the delta epoch given by Δ. This is used to update the rates of accumulator on a trial by trial basis. b. Evolution of the action choice rates for each symbol-action learning. c. The profile of neural activity in the delta epoch with learning from experimental data (left) and the model (right). d. Learning curves of each symbol-action association learning from experimental data (left) and the model (right). e. Strategy used by the monkey during learning (left) and the model (right).

**Supplementary Table 1:** Distribution of P-cells with respect to hand movement selectivity and delta epoch

|  |  |
| --- | --- |
| Total number of P-cells | 128 |
| Hand movement selective | 106 |
| P-cells with a delta epoch | 105 |
| Hand movement selective but with no delta epoch | 18 |
| Hand movement non-selective but did have a delta epoch | 17 |
| Neither hand movement selective, nor with a delta epoch | 5 |

**Supplementary Table 2:**

| Alternate hypothesis | Rebutting data | Explanation |
| --- | --- | --- |
| P-cells encoded specific motor kinetics | Hand: Fig 2F-J S1C, S10 H-J;<br>Eye: compare Figs S10K-M with 3C-F<br>Lick: compare Figs S10N with 3C-F;<br>also see S5 B-C | These P-cells encoded gross hand-movement and not specific hand-movement kinetics. Nor did they encode other movements including eye movements and licking. |
| P-cells encoded specific sensory features | Fig S1B S10A-B | These P-cells were not tuned to specific visual stimuli |
| P-cells encoded reward information during the performance of an overtrained task | Fig 3G-H | P-cells encoded reward information only when it was relevant to visuomotor association learning |
| P-cells encoded reward expectation signal | Fig S5 | The increase in activity before reward delivery was a task-related motor signal and was not a reward expectation signal |
| P-cells changed activity at the visuomotor symbol switch due to changes in motor kinematics (eye, hand, licking etc) | Hand: Fig 2F-G<br>Eye: compare Figs S10K-M and 3C-F<br>Lick: compare Figs S10N with 3C-F;<br>also see S5 B-C | Firstly, P-cells encoded specific motor kinetics. Second, hand movement kinetics, licking, eye movements remained similar. |
| P-cells changed activity at the visuomotor symbol switch due to changes in sensory features | Fig S9 | P-cells changed activity at the visuomotor symbol switch due to changes in 'learning'. They changed their activity at a switch even when the sensory features remained the same (reversal learning) |
| P-cells changed activity at the visuomotor symbol switch due to changes in reaction time | Fig S4 | P-cells also changed activity at the visuomotor symbol switch when there were no changes in reaction time |
| Delta epoch was purely due to chance | Fig S8 | Delta epoch was not purely due to chance or threshold crossing. |
| Delta epoch was due to different sensory stimuli shown | Fig S10 A-B | Activity in delta epoch was similar for either stimulus |
| Delta epoch was due to the choice of different hands | Fig S10 C-D | Activity in delta epoch was similar for either hand |
| Delta epoch was due to changes in eye position | compare Figs S10K-M with 3C-F | Monkey were free to move their eyes. Eye position did not correlate with neural activity |
| Delta epoch was due to changes in licking behavior | compare Figs S10N with 3C-F; also see S5 B-C | Licking did not correlate with neural activity |
| Delta epoch was due to differences in solenoid clicks | Fig 3G-H | Solenoid clicks were conditionally paired with reward delivery. Neural activity was correlated with reward delivery only during learning |
| Delta epoch was due to differences in reward delivery | Fig 3I-J | Different amount of reward delivery did not affect neural activity |
| Delta epoch was due to differences in correct vs wrong hand movement of the hand that was used to report the choice | Fig S10 K | The hand movements after correct and wrong trials for the released hand were similar. |
| Delta epoch was due to differences in correct vs wrong hand movement of the hand that was not used to report the choice | Fig S10 I | The hand movements after correct and wrong trials for the released hand that was not were similar. |
| Delta epoch was due to differences in the time of return of correct vs wrong hand movement | Fig S10 J | The time of return of correct vs wrong hand movement were similar. |
| Delta epoch encoded the memory of history of decisions before the most recent one and predicted the impending decision | Fig 3A-B, 4F-H, Fig S6, S7 | Activity in the delta epoch reflected a memory of the outcome of single, most recent decision |

## References and Notes

1. M. Ito, The Cerebellum and Neural Control. Control (Raven Pr, 1984).

2. Thach, S. G. L. W. T. in Principles of Neural Science Vol. 5 (ed Eric R. Kandel; James H. Schwartz; Thomas M. Jessell; Steven A. Siegelbaum; A. J. Hudspeth) Ch. 42, 960–981 (McGraw-Hill Education / Medical, 2012)

3. J. L. Raymond, S. G. Lisberger, M., The cerebellum: a neuronal learning machine? Science (1996).

4. J. C. Eccles, M. Ito, J. Szentagothai, The Cerebellum as a Neuronal Machine. (Springer Berlag, Berlin, 1967).

5. M. Manto et al., Consensus paper: roles of the cerebellum in motor control—the diversity of ideas on cerebellar involvement in movement. The Cerebellum 11, 457–487 (2012).

6. F. Mjp, Recherches expérimentales sur les propriétés et les fonctions du système nerveux, dans les animaux vertébrés (ed 1). (Chez Crevot, Paris, 1824), vol. vol 26.

7. J. F. Medina, S. G. Lisberger, Links from complex spikes to local plasticity and motor learning in the cerebellum of awake-behaving monkeys. Nature neuroscience 11, 1185–1192 (2008).

8. S. G. Lisberger, The neural basis for learning of simple motor skills. Science 242, 728–735 (1988).

9. J. S. Albus, A theory of cerebellar function. Mathematical Biosciences 10, 25–61 (1971).

10. D. Marr, A theory of cerebellar cortex. The Journal of Physiology 202, 437–470 (1969).

11. C. L. Ojakangas, T. J. Ebner, Purkinje cell complex and simple spike changes during a voluntary arm movement learning task in the monkey. Journal of neurophysiology 68, 2222–2236 (1992).

12. T. J. Ebner, A. L. Hewitt, L. S. Popa, What features of limb movements are encoded in the discharge of cerebellar neurons? The Cerebellum 10, 683–693 (2011).

13. A. V. Roitman, S. Pasalar, M. T. Johnson, T. J. Ebner, Position, direction of movement, and speed tuning of cerebellar Purkinje cells during circular manual tracking in monkey. Journal of Neuroscience 25, 9244–9257 (2005).

14. C. J. Stoodley, J. P. MacMore, N. Makris, J. C. Sherman, J. D. Schmahmann, Location of lesion determines motor vs. cognitive consequences in patients with cerebellar stroke. NeuroImage: Clinical 12, 765–775 (2016).

15. J. D. Schmahmann, Dysmetria of thought: clinical consequences of cerebellar dysfunction on cognition and affect. Trends in Cognitive Sciences 2, 362–371 (1998).

16. E. B. Becker, C. J. Stoodley, in International review of neurobiology. (Elsevier, 2013), vol. 113, pp. 1–34.

17. G. Zhao, K. Walsh, J. Long, W. Gui, K. Denisova, Reduced structural complexity of the right cerebellar cortex in male children with autism spectrum disorder. PloS one 13, e0196964 (2018).

18. J. A. Mangels, R. B. Ivry, N. Shimizu, Dissociable contributions of the prefrontal and neocerebellar cortex to time perception. Cognitive Brain Research 7, 15–39 (1998).

19. R. L. Buckner, The cerebellum and cognitive function: 25 years of insight from anatomy and neuroimaging. Neuron 80, 807–815 (2013).

20. P. L. Strick, R. P. Dum, J. A. Fiez, Cerebellum and nonmotor function. Annual review of neuroscience 32, 413–434 (2009).

21. R. M. Kelly, P. L. Strick, Cerebellar loops with motor cortex and prefrontal cortex of a nonhuman primate. The Journal of neuroscience : the official journal of the Society for Neuroscience 23, 8432–8444 (2003).

22. D. Caligiore et al., Consensus Paper: Towards a Systems-Level View of Cerebellar Function: the Interplay Between Cerebellum, Basal Ganglia, and Cortex. Cerebellum (London, England) 16, 203–229 (2017).

23. A. C. Bostan, R. P. Dum, P. L. Strick, Cerebellar networks with the cerebral cortex and basal ganglia. Trends in cognitive sciences 17, 241–254 (2013).

24. H. C. Leiner, A. L. Leiner, R. S. Dow, Does the cerebellum contribute to mental skills? Behavioral neuroscience 100, 443–454 (1986).

25. L. S. Popa, A. L. Hewitt, T. J. Ebner, The cerebellum for jocks and nerds alike. Frontiers in Systems Neuroscience 8, (2014).

26. L.-H. C. review, Solving the mystery of the human cerebellum. Neuropsychology review, (2010).

27. S. Tomatsu et al., Information processing in the hemisphere of the cerebellar cortex for control of wrist movement. Journal of neurophysiology 115, 255–270 (2015).

28. M. J. Wagner, T. Kim, J. Savall, M. J. Schnitzer, L. Luo, Cerebellar granule cells encode the expectation of reward. Nature 544, 96–100 (2017).

29. D. O. Kellett, I. Fukunaga, E. Chen-Kubota, P. Dean, C. H. Yeo, Memory consolidation in the cerebellar cortex. PLoS One 5, (2010).

30. W. Heffley et al., Coordinated cerebellar climbing fiber activity signals learned sensorimotor predictions. Nature neuroscience 21, 1431–1441 (2018).

31. F. A. Miles, S. G. Lisberger, Plasticity in the vestibulo-ocular reflex: a new hypothesis. Annual review of neuroscience 4, 273–299 (1981).

32. L. S. Popa, M. L. Streng, A. L. Hewitt, E.-T. J. Cerebellum, The errors of our ways: understanding error representations in cerebellar-dependent motor learning. The Cerebellum, (2016).

33. J. L. Raymond, Lisberger.-S. G. of Neuroscience, Neural learning rules for the vestibulo- ocular reflex. Journal of Neuroscience, (1998).

34. Y. Yang, S. G. Lisberger, Purkinje-cell plasticity and cerebellar motor learning are graded by complex-spike duration. Nature 510, 529–532 (2014).

35. R. S. Sutton, A. G. Barto, Introduction to reinforcement learning. (MIT press Cambridge, 1998), vol. 135.

36. A. L. Person, Raman. -I. M Purkinje neuron synchrony elicits time-locked spiking in the cerebellar nuclei. Nature, (2012).

37. A. V. Hays, B. J. Richmond, L. M. Optican, A UNIX-based multiple process system for real-time data acquisition and control. WESCON Conf. Proc. 2, 1–10 (1982).

38. G. Dijck et al., Probabilistic identification of cerebellar cortical neurones across species. PloS one 8, (2013).

39. J.-Y. Y. Tinevez et al., TrackMate: An open and extensible platform for single-particle tracking. Methods (San Diego, Calif.) 115, 80–90 (2017).

40. J. Schindelin et al., Fiji: an open-source platform for biological-image analysis. Nature methods 9, 676–682 (2012).

41. K. Jaqaman et al., Robust single-particle tracking in live-cell time-lapse sequences. Nature methods 5, 695–702 (2008).

42. S. Fusi, W. F. Asaad, E. K. Miller, W.-X. J. Neuron, A neural circuit model of flexible sensorimotor mapping: learning and forgetting on multiple timescales. Neuron, (2007).

43. S. Jana, A. Gopal, A. Murthy, Evidence of common and separate eye and hand accumulators underlying flexible eye-hand coordination. American Journal of Physiology-Heart and Circulatory Physiology, (2016).

